# RSV competes with the host for translational machinery without a host shutoff strategy

**DOI:** 10.64898/2026.08.31.748072

**Authors:** Kyra Kerkhofs, Nicholas R. Guydosh

**Affiliations:** Laboratory of Biochemistry and Genetics, National Institute of Diabetes and Digestive and Kidney Diseases, National Institutes of Health, Bethesda, MD, 20892

**Keywords:** RSV, translation efficiency, host shutoff, dsRNA sensors, ribosome profiling, integrated stress response, eIF4E

## Abstract

RNA viruses often enhance ribosome recruitment to their own mRNAs through non-canonical sequence elements or by degrading host mRNA. Respiratory syncytial virus (RSV) produces mRNAs with host-like features, including 5’-cap and poly(A) tail. Therefore, the virus lacks an obvious mechanism to preferentially protect its own mRNAs or recruit ribosomes. Furthermore, it remains unknown how RSV interacts with antiviral defense pathways that would reduce cap-dependent translation. Using spike-in normalized sequencing of total and ribosome-associated RNA, we found that RSV does not appear to evoke any host shutoff mechanisms to limit the expression of host genes. These findings show that RSV manages to make use of available ribosomes by competing effectively with host mRNAs and any translational shutoff mechanism would be detrimental. Consistent with this, we found that following activation of antiviral host pathways that reduce cap-dependent translation, translation of RSV mRNAs is decreased to the same extent as host mRNAs. Furthermore, we found that RSV infection does not trigger the dsRNA-activated kinase PKR (which initiates the ISR) and OAS (activates endonuclease RNase L) pathways. These data support a model in which RSV achieves viral protein production, not though inhibiting the host, but by successfully competing with host mRNAs and avoiding activation of antiviral pathways.

**Highlights:**

- RSV does not induce host mRNA degradation
- RSV mRNAs effectively recruit ribosomes throughout infection and cause some reduction in host mRNA translation
- RSV mRNAs depend on eIF4E for translation and are stabilized independently of LARP1 when eIF4E is depleted
- RSV avoids strong activation of the ISR and OAS/RNaseL pathways instead of directly inhibiting their activation

## Introduction

Most viruses are fully dependent on the host translational machinery to produce their viral proteins (Walsh and Mohr 2011; Fels et al. 2026). Viral mRNAs therefore are required to compete against abundant host mRNAs for ribosomes (Gale et al. 2000). A common strategy employed by viruses to win this competition is to induce “host shutoff” and therefore reduce host gene expression and increase the availability of ribosomes and associated machinery for viral protein production. Another advantage of host shutoff is that it strongly reduces the antiviral innate immune response. Host shutoff often involves reducing host mRNA levels or restricting access of host mRNAs to ribosomes (Abernathy and Glaunsinger 2015; Stern-Ginossar et al. 2019). Interestingly, host cells are equipped with antiviral pathways that also effectively shut off translation, resulting in neither host nor virus being able to express genes (McCormick and Khaperskyy 2017). Therefore, viruses need to both access the limited translational machinery and simultaneously prevent activation of these antiviral defense pathways. Respiratory syncytial virus (RSV) is a single-stranded negative-sense RNA virus that produces ten individual mRNAs that mimic important host mRNA features, including a 5′-cap, poly(A)-tail and untranslated regions (UTRs) flanking the main open reading frame (ORF) (Cao et al. 2021; Donovan-Banfield et al. 2022). Viral transcription and replication takes place in liquid-liquid phase separated inclusion bodies within the cytoplasm. Following transcription, viral mRNAs are exported into the cytoplasm for translation (Rincheval et al. 2017). Our recent work showed that RSV infection increases ribosome loading and remodels the host translatome (Kerkhofs et al. 2025). However, limitations in the ribosome sedimentation methodology used there left open questions about what portion of translating mRNAs are viral and how the recruitment of ribosomes to viral mRNA impacts host mRNA translation.

Cap-dependent translation initiation starts with assembly of the preinitiation complex, which contains the 40S ribosomal subunit, initiator methionine tRNA and eIF2 protein. Next, the preinitiation complex is recruited to the 5′-end of the mRNA by cap-binding complex eIF4F, followed by scanning and start codon selection. The cap-binding complex consists of the helicase eIF4A, cap-binding protein eIF4E and scaffold protein eIF4G (Hinnebusch 2014). Two main cellular translation regulation pathways that can limit translation initiation include reduced mTOR signaling and the integrated stress response (ISR) (Walsh et al. 2013). Reduced mTOR signaling activates the eIF4E-binding proteins (4E-BPs), resulting in reduced availability eIF4E for eIF4F assembly (Ma and Blenis 2009). In addition, under conditions where mTOR signaling is reduced, LARP1 associates with 5′-TOP mRNAs and suppresses their translation (Fonseca et al. 2015). The ISR functions by effectively limiting eIF2 availability by phosphorylation of the α-subunit, leading to a global reduction in protein synthesis (Pakos-Zebrucka et al. 2016). An important negative feedback loop associated with the ISR operates through the protein GADD34, the regulatory subunit of the phosphatase complex which dephosphorylates eIF2α-P (Novoa et al. 2001). However, during HIV-1 infection, GADD34 functions as an antiviral protein by reducing viral translation through interactions made by its protein domains that are independent of its phosphatase function (Ishaq et al. 2020). GADD34 can be transcriptionally activated by the ISR (Ma and Hendershot 2003) and by interferon (IFN) (Ruggieri et al. 2012) and, unlike most mRNAs, translationally activated by eIF2α phosphorylation (Lee et al. 2009). Interestingly, many viral mRNAs containing internal ribosomal entry sites (IRES) have been shown to bypass the dependency on eIF2 and eIF4E and thus maintain viral protein production following activation of stress-induced pathways (Kwan and Thompson 2019).

A hallmark of viral infection is the appearance of dsRNA in the cell, which is detected by several receptors that activate the innate immune system to help limit the virus. Activation of RIG-I-like receptors (RLRs) leads to production of IFN, which in turn induces the expression of IFN-stimulated genes (ISGs) to regulate innate immune sensing and restrict viral replication (Rehwinkel and Gack 2020). Another dsRNA sensor is protein kinase R (PKR), an activator of the ISR that leads to inhibition of all cap-dependent translation (Cesaro and Michiels 2021). A third dsRNA receptor, oligoadenylate synthase (OAS), synthesizes 2’–5’-oligoadenylate (2–5A) which, in turn, activates RNase L (Drappier and Michiels 2015). RNase L is an endonuclease that targets both host and viral RNA at UN^N motifs (Han et al. 2014; Karasik and Guydosh 2024). It remains an open question if RSV manages to efficiently produce viral proteins because it escapes outcomes of dsRNA sensing or if it avoids activation of these pathways.

It remains to be understood how RSV successfully competes with host mRNAs for the translational machinery while limiting activation of antiviral pathways. Previous work reported that while RSV infection activates PKR, the downstream phosphorylation of eIF2α remains limited through blocking the binding of PKR to eIF2α and by enhancing phosphatase activity (Groskreutz et al. 2010; Lindquist et al. 2010a). Antiviral immune signaling was shown to be dampened by RSV-NS1 and -NS2 proteins which block immune receptors and downstream signaling (Pei et al. 2021; Lo et al. 2005; Thornhill and Verhoeven 2020). Inclusion bodies were shown to concentrate genomes, antigenomes and mRNAs produced by RSV, which facilitates evasion of innate immune receptors which are generally excluded from these sites (Hwang et al. 2026; Rincheval et al. 2017). Overall, these studies suggest that RSV possesses multiple antagonistic mechanisms to reduce host-induced antiviral pathways.

In this study we asked how RSV competes for limiting translational machinery and interacts with antiviral defense pathways that can reduce global cap-dependent translation. Using spike-in normalized RNA sequencing, we present evidence that RSV efficiently recruits host ribosomes but does not actively reduce host mRNA levels. We therefore conclude that RSV can effectively express its genes by producing abundant transcripts and recruiting ribosomes efficiently enough to obtain about 30% of ribosomes in the cell. We also demonstrate that activation of the antiviral host pathways that reduce cap-dependent translation affect RSV mRNAs to a similar extent as host mRNAs. Consistent with this, RSV infection is not associated with a strong ISR induction nor activation of RNase L. We therefore propose that RSV maintains cap-dependent translation via a “fair competition” model where host and virus each recruit a fraction of the ribosome population to their mRNAs to carry out translation.

## Results

### RSV infection does not induce host mRNA degradation

We established a system for studying RSV infection where we infected A549 lung carcinoma cells with virus and harvested cells over the course of infection at 6, 12, 18 and 24 hours (see Methods) (**Figure 1A**). We infected cells with an MOI of 3 and confirmed this was a sufficient level to infect the majority of cells (**Figure S1A**) and induce a steady increase in viral protein as infection progressed (**Figure S1B**). We extracted total RNA from these cells and added an identical amount of ERCC spike-in to each sample that had been prepared to contain the same amount of total RNA (see Methods) (**Figure 1A**). Following high-throughput sequencing, we observed a similar number of ERCC spike-in reads in the different samples, consistent with little change in overall mRNA levels in the cell (**Supplementary Figure 1C**).

**Figure 1.**
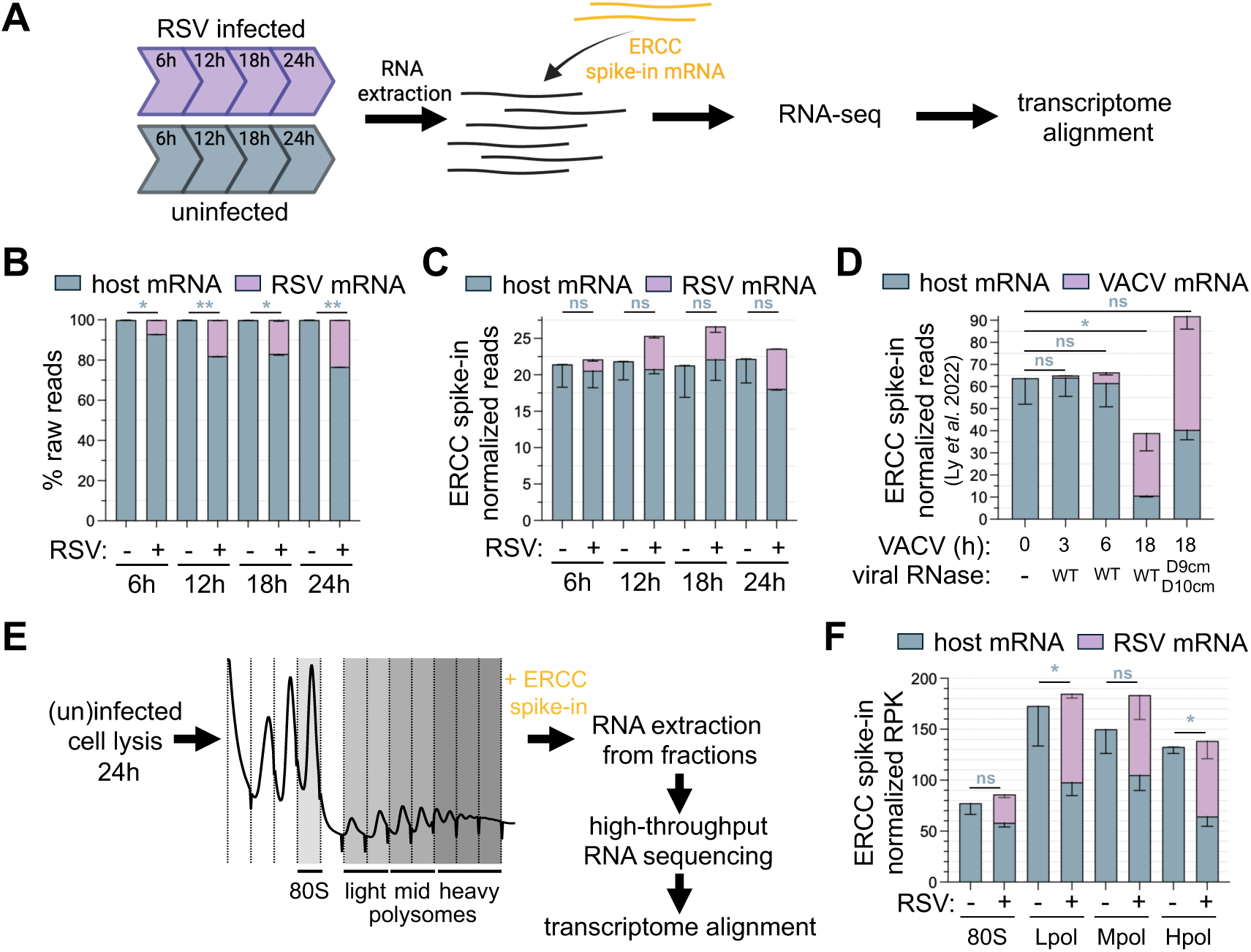
RSV mRNAs compete with host mRNA for ribosomes without causing host shutoff. (A) Schematic of the ERCC spike-in normalized RNA-seq experiment for RSV infection (MOI 3) timecourse. (B,C) Quantification of the abundance of host and viral mRNAs, shown normalized to total raw read counts (B) and ERCC spike-in read abundance (C). A paired t-test was performed to compare host mRNA abundance (blue bars) between uninfected and infected conditions for each timepoint separately. Spike-in normalization reveals that these is no significant loss of absolute levels of host mRNA during RSV infection. Bars represent the mean and error bars indicate the standard deviation. (D) VACV ERCC spike-in normalized RNA-seq dataset downloaded from SRA (SRP340405), showing absolute host and viral mRNA abundance. VACV D9cm D10cm are catalytic mutants (cm) of the viral decapping enzymes. A paired t-test was performed to compare host mRNA abundance between uninfected (0 hpi) and infected cells at each timepoint. VACV induces a strong reduction in host mRNA after 18 hours. Bars represent the mean and error bars indicate the standard deviation. (E) Schematic of the ERCC spike-in containing sucrose gradient fractionation RNA-seq experiment following RSV infection (MOI 3, 24h). (F) Quantification of ERCC spike-in normalized reads, also normalized by mRNA length (per kilobase, RPK), for host and virus within each fraction of the gradient. A paired t-test was performed to compare host mRNA abundance between uninfected and infected fractions. RSV infection induces a slight decrease in host mRNA in some polysome fractions. 80S: monosome fraction, Lpol: light polysomes, Mpol: middle polysomes and Hpol: heavy polysomes. Bars represent the mean and error bars indicate the standard deviation. P values < 0.01 are indicated by two asterisks (**), P values < 0.05 are indicated by one asterisks (*) and ns indicates not significant.

We then asked whether RSV infection induces host mRNA degradation. Analysis of the RNA-seq data without taking spike-in normalization into account showed an apparent steady decrease in host mRNA as the infection progressed. This significant decrease was proportionately compensated by an increase in viral mRNA levels, as expected since RNA levels without spike-in normalization are relative (**Figure 1B**, blue bars). In contrast, the information from spike-in normalization shows how the absolute amount of mRNA in the cell changes during an experiment (Burke et al. 2019; Ly et al. 2022; Chen et al. 2016). Interestingly, the analysis with ERCC spike-in normalization showed that RSV did not induce significant loss of absolute levels of host mRNA (**Figure 1C**). Consistent with this observation, cells appeared to accumulate slightly higher amounts of total mRNA during infection though this did not reach a significant threshold (**Figure 1C**, the sum of host and viral mRNA). For comparison, we performed a similar analysis on a dataset utilizing ERCC spike-ins from cells infected with vaccinia virus (VACV) (Ly et al. 2022). These data serve as a positive control for the absolute loss of host mRNA since VACV induces mRNA degradation through expression of two viral decapping enzymes, D9 and D10 (Parrish and Moss 2007; Parrish et al. 2007). As expected (Ly et al. 2022), absolute host mRNA levels were strongly reduced after 18h of VACV infection (**Figure 1D**, compare WT timecourse). This loss was dampened in data from a virus lacking both viral decapping enzymes (D9 and D10) (**Figure 1D**, compare VACV deletion mutant against WT at 18 hpi).

Although absolute host mRNA levels do not change much after RSV infection, individual host mRNAs can undergo significant changes. Therefore, we measured the differential expression of host mRNA levels with ERCC normalization in DESeq2 (**Table S1**). We observed a strong interferon (IFN) response throughout the RSV infection timecourse (**Figure S1D**, RSV), as previously reported (Kerkhofs et al. 2025). In contrast, VACV dampens the IFN response, and this was dependent on the decapping enzymes D9 and D10 (**Table S2**, **Figure S1D**, VACV). The downregulated IFN transcripts are the receptor genes which are constitutively expressed (**Figure S1D**, dark green squares), while the downregulated ISG transcripts included transcripts with basal levels of expression that were strongly reduced following VACV infection (**Figure S1D**, light green circles). Overall, our data show that RSV does not cause substantial degradation of host mRNA, either directly with its own proteins or by activating host decay pathways such as OAS/RNase L, and this leaves the host free to activate transcriptional programs in response.

### Translation capacity is slightly increased during RSV infection

Since RSV does not degrade host mRNAs, its mRNAs must compete with those of the host for ribosomes. We next investigated whether RSV utilizes any strategies to bias this competition in its favor by inhibiting host mRNA translation. An established example of a virus that employs this strategy is poliovirus, which expresses a viral protease that specifically cleaves the essential 5′ cap-binding translation initiation factor eIF4G in order to globally reduce host protein production (Gradi et al. 1998). The mRNA of poliovirus bypasses this restriction through use of an internal ribosome entry site (IRES) in its 5′ UTR, which initiates translation independent of the 5′ cap (Pelletier and Sonenberg 1988). In our previous work, we found that RSV induces heavier polysomes during infection by polysome profiling (Kerkhofs et al. 2025), a method which separates monosomes and polysomes by density centrifugation. While this result suggests higher engagement of ribosomes with mRNA during infection, it remains to be determined what fraction of the ribosomes are occupied by viral versus host mRNAs. For example, polysomes formed in cells infected with poliovirus are mostly occupied by viral mRNAs since host mRNAs rely on eIF4G for translation initiation (Cardinali et al. 1999).

To determine the absolute number of viral mRNAs associated with ribosomes, we performed RNA-seq on different polysome species obtained 24 hours after infection and compared these to a matching uninfected control (see Methods) (**Figure 1E**). In particular, we examined the 80S monosome fraction and three polysome species that we define as light polysomes (Lpol, 2-3 ribosomes), middle polysomes (Mpol, 4-5 ribosomes) and heavy polysomes (Hpol, 6+ ribosomes). For this analysis, ERCC spike-ins were used to normalize for RNA extraction efficiency and to allow for quantification of absolute mRNA levels within each of the four fractions. In addition, to account for differences in gene length, we visualized the data as RPK (reads per kilobase). We observed a moderate decrease in host mRNAs across all four ribosome-bound fractions (**Figure 1F**, blue bars) and these were replaced by viral mRNAs. Interestingly, there was a slight increase in the total (host and viral) number of mRNAs that are associated with ribosomes, which is consistent with our previous finding of increased polysomes during RSV infection (Kerkhofs et al. 2025).

To confirm the translation levels measurements from **Figure 1F** with a related method, we determined global translation levels by using the SUrface SEnsing of Translation (SUnSET) assay, which measures puromycin incorporation (during elongation, prior to termination) into the nascent polypeptide chain. Nascent proteins were detected by using a puromycin-specific antibody. We confirmed the functionality of the assay with pre-treatment with cycloheximide or arsenite, which inhibit translation (**Supplementary Figure 2A**). Next, we determined puromycylation levels between mock- and RSV-infected conditions after 24 hours by western blot and found that overall translation levels (combined total of both host and virus nascent chains) remain similar during infection (**Supplementary Figure 2B**).

Overall, these data are consistent with a moderate reduction in translation of host mRNAs that is compensated by an increase in translation of viral mRNAs. However, whether the reduction in host translation is caused by RSV actively blocking the ability of host mRNAs to recruit ribosomes or viral mRNAs simply being effective at recruiting ribosomes from the limited pool more effectively remains to be determined.

### Total rRNA levels remain unchanged during infection

Consistent with our previous findings that showed some increase in heavier polysomes during RSV infection (Kerkhofs et al. 2025), we observed somewhat higher numbers of all mRNAs (sum of host and virus) associated with polysomes when comparing infected against uninfected cells (**Figure 1F**). These data suggest that RSV may expand the translational capacity in the cell. Higher translational capacity could be caused by an increase in number of ribosomes in the cell through augmented ribosomal RNA (rRNA) transcription and ribosomal protein production (*i.e.* ribosome biogenesis). To test whether rRNA levels increased, we determined the cell count, performed in-plate lysis of cells with spike-in RNA containing Trizol (to serve as a normalization for extraction efficiency in each sample), extracted RNA from the samples, and then measured rRNA levels with tapestation analysis (**Supplementary Figure 2C**). As a control for reduced ribosome levels, we stressed cells through Torin1 treatment or serum starvation for 24 hours. These data showed a strong decrease in rRNA levels (**Supplementary Figure 2C**) that was correlated with a reduced cell count. This indicates that rRNA per cell remained about the same as cells likely divided less. In the case of RSV infection, rRNA levels and cell count remained about the same, indicating that cells continued to divide and that RSV did not increase rRNA levels to enhance translational capacity (**Supplementary Figure 2C**).

### Ribo-seq confirms viral mRNAs recruit a substantial fraction of translating ribosomes

We showed that viral mRNAs recruit a considerable fraction of ribosomes during infection (see **Figure 1E,F**). To more quantitatively and precisely compute how ribosomes distribute between host and viral transcripts, we performed ribosome profiling (ribo-seq). This method uses RNase digestion of cell lysates to produce ribosome-protected footprints (**Figure 2A**). The length distribution of the ribosome-protected fragments was in the expected 28-30 nt range for both host and virus, including the slightly longer footprint known to be specific to stop codons (Kerkhofs and Guydosh 2026; Ingolia et al. 2011) (**Figure S3A**). Biological replicates of mRNA-Seq (same data as in **Figure 1**) and ribo-seq (prepared from the same lysates) grouped together in a principal component analysis (PCA) plot, indicating high reproducibility (**Figure S3B**, ribo-seq). As expected, there was a clear separation between the RNA-seq and ribo-seq datasets (**Supplementary Figure 3B**, PC1, compare circles and triangles). In addition, we observed a clear separation between mock- and RSV-infected samples, with stronger effects as the timecourse advanced, presumably reflecting the increasing activation of the innate immune response (**Supplementary Figure 3B**, PC2). Analysis of ribosome footprint distribution between host and virus revealed that RSV obtains about 30% of translating ribosomes after 24 hours of infection with a gradual increase during earlier timepoints (**Figure 2B**, ribo-seq). This indicates that RSV is not completely overtaking the translational machinery, consistent with our results in **Figure 1**. Viral ribo-seq and RNA-seq levels increased at similar rates over time (**Figure 2B**), as expected for constant levels of viral translation over time.

**Figure 2.**
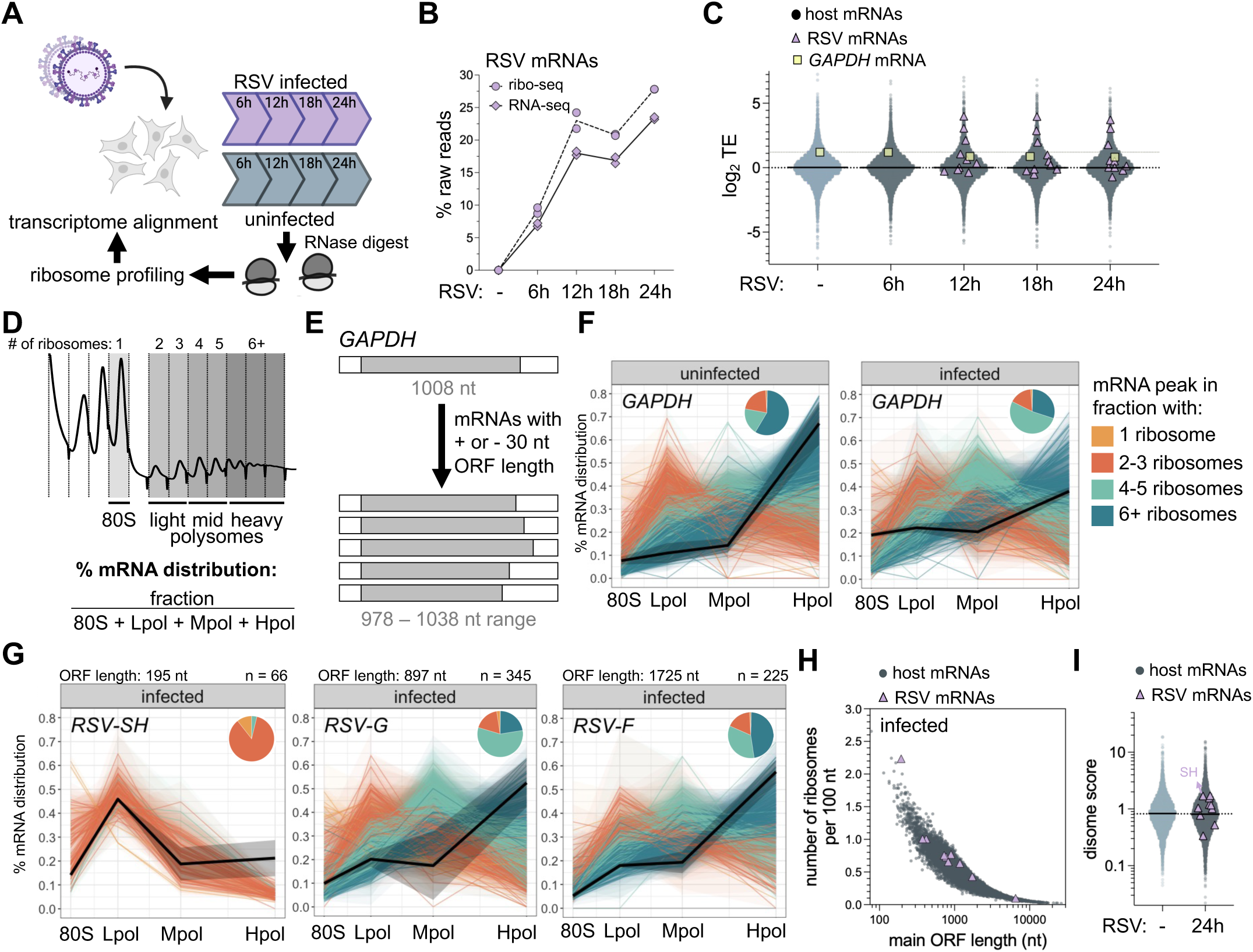
RSV mRNAs are translated as efficiently as well-translated host mRNAs. (A) Schematic of the ribo-seq experimental workflow for an RSV infection (MOI 3) timecourse, matching the RNA-seq timecourse datasets in Figure 1. (B) Quantification of the relative abundance of viral mRNAs over time, shown as a percentage of raw read counts for ribo-seq and RNA-seq. RSV mRNAs maintain a constant level of translation over time and occupy almost 30% of the ribosomes after 24 hours. Each datapoint indicates the mean of all ten viral mRNAs from a biological replicate. (C) TEs calculated by DESeq2 for all individual mRNAs. All ten RSV mRNAs and GAPDH are highlighted. The black dotted line indicates the average TE across all transcripts and the grey dotted line the TE of GAPHD in uninfected cells. (D) Calculation of mRNA distribution across the gradient. The percentage in each fraction was calculated by dividing the mRNA abundance in each fraction by the total abundance across all fractions, with the sum of all fractions adding up to 100%. (E) Schematic representation of selection of transcripts with similar ORF length as the highlighted transcript of interest (*GAPDH* in the example). (F,G) Line graphs showing the distribution of *GAPDH* (F) and individual viral mRNAs (G) across the sucrose gradient. The black line represents the average of the transcript of interest, and standard deviation is shown as shaded areas. Host mRNAs with similar ORF lengths (±30 nt) are colored according to their most abundant location within the gradient (80S: light orange, Lpol: dark orange, Mpol: green, Hpol: blue). Pie charts show the proportion of dominant fraction of each transcript across the gradient. The ORF length of the viral transcripts is indicated above as well as the number (n) of host transcripts included in the analysis. 80S: monosome fraction, Lpol: light polysomes, Mpol: middle polysomes and Hpol: heavy polysomes. (H) Ribosome density of individual host and RSV mRNAs during infection, calculated as number of ribosomes per 100 nt ORF length. Graph demonstrates that the viral RNAs have ribosome densities that are similar to well-translated host RNAs and that, overall, shorter ORFs tend to have higher ribosome density (as expected). (I) The disome score for host and RSV mRNAs defined as the ratio between ORF reads from disome-seq and matching monosome-seq.

### Viral transcripts are translated as efficiently as well-translated host transcripts

Next, we determined the translation efficiency (TE) of viral mRNAs over time by using DESeq2 to calculate the ratio of ribo-seq over RNA-seq counts (**Table S3**). TE serves as a measure of how efficiently individual mRNAs are loaded with ribosomes. Generally, the TE of viral transcripts fell within a range comparable to that of host transcripts (**Figure 2C**, purple triangles), indicating that viral mRNAs are efficiently translated, and this property remained constant over time.

To confirm this finding, we also developed a method to analyze the distribution of each mRNA across our sucrose gradient datasets (see **Figure 1E,F**). Since we added spike-in to each of the four fractions taken from the sucrose gradient, differences in RNA extraction efficiency between fractions are taken into account and this facilitates comparisons between the fractions. For each mRNA, we summed up the ERCC spike-in normalized read count across all fractions and calculated the fraction of this total in each fraction (%mRNA distribution) (**Figure 2D, Table S4**). For example, the host mRNA *GAPDH* main ORF is 1008 nt in length, which provides enough space to accommodate many ribosomes, and is known to be well translated. These features are consistent with its high abundance in the heavy polysomes of uninfected cells. (**Figure 2E,F**, black line, uninfected). Since the number of ribosomes found on a given mRNA depends on the mRNA length, we only compared transcripts with similar (±30 nt, approximate footprint of a ribosome) main ORF lengths (**Figure 2E**). Under uninfected conditions, transcripts with a length similar to *GAPDH* were mostly associated with heavy polysomes (**Figure 2F**, dark green slice in pie chart). Interestingly, following RSV infection, *GADPH* (and mRNAs of similar length) moved from the heavy polysomes into lighter fractions (**Figure 2F**, compare lines and pie charts). This shift of *GAPDH* into lighter polysome fractions is also reflected in the ribo-seq TE timecourse (see **Figure 2C**, squares move below the dotted line). Consistent with the analysis of **Figure 1F**, a length-sorted analysis of all host mRNAs also revealed a global shift of host mRNAs out of heavy polysomes following infection (**Figure S4**, **Table S4**).

Next, we applied this length-constant analysis to the abundance of viral mRNAs within the sucrose gradient. First, we investigated viral envelope proteins SH, G and F, which have ORF lengths of 195, 897 and 1725 nt, respectively. We found that the shortest ORF transcript, RSV-SH, is found predominantly in the light polysome peak (>40%), comparable to host mRNAs of similar length (**Figure 2G**). However, about 40% of the SH mRNA is found in heavier fractions, consistent with it being well translated with potential implications in ribosome collisions (discussed later). Next, RSV-G and -F were predominantly found in the heavy polysome fraction (>50%). This is expected for the RSV-F transcript, which contains a long ORF. However, RSV-G contains a shorter ORF with less than 25% of host transcripts predominantly found in a similar heavy polysome fraction (**Figure 2G**), suggesting it is also particularly well translated. Much like RSV-F, the other viral mRNAs distributed similarly to host mRNAs with similar ORF length (**Figure S5A**).

Next, we defined an apparent ribosome density for each host and viral mRNA by first multiplying each of the four %mRNA distribution number by the expected number of ribosomes found within that fraction (**Table S5**). For example, the monosome fractions were multiplied by 1, while the light polysomes by 2.5 (since this fraction contains mRNAs with 2 or 3 ribosomes loaded) (see Methods) (see **Figure 2D**). We then divided this number by the ORF length to give an approximation of the density of ribosomes within the open-reading frame (ribosomes per 100 nt, listed in **Table S5**). Plotting ribosome density for mRNAs against ORF length showed a trend where host transcripts with short ORFs had higher ribosome density compared to those with longer ORFs (**Figure 2H**, **Figure S5B**), as shown previously (Arava et al. 2003; Thompson and Gilbert 2017). Viral mRNAs generally fell within the expected ribosome density range for host mRNAs (**Figure 2H**), consistent with the earlier analysis using the ribo-seq method (see **Figure 2C**).

### Viral mRNAs exhibit similar ribosome collision rates as host mRNAs

Given that the main ORF length of SH is less than 200 nt, the observation that some transcripts are in the heavy polysome fraction (6+ ribosomes) suggests the mRNA is heavily loaded with ribosomes, which would lead to a higher probability of ribosome collisions occurring. Since an elongating ribosome occupies around 30 nt, in the most extreme case a fully loaded RSV-SH mRNA could fit about 7 ribosomes. We confirmed that host mRNAs with short ORFs (< 500 nt) are not abundantly found in the heavy polysomes (**Figure S4**). However, we noted in **Figure 2G** that around 40% of the RSV-SH mRNA is found in middle and heavy polysome fractions, which are loaded with 4 ribosomes or more. Therefore, we tested if translation of RSV-SH mRNA results in a large number of ribosome collisions (also referred to as disomes). For this, we performed disome profiling (Meydan and Guydosh 2020; Marks et al. 2026) and determined a disome score by calculating the ratio between reads derived from disome-seq (TPM for footprint sizes between 50-70 nt) and matching monosome-seq (TPM for footprint sizes between 25-34 nt) for translated transcripts (basemean > 100) (**Table S6**). Overall, all RSV mRNAs fell within the host mRNA disome score range, indicating a similar level of ribosome collisions on these transcripts (**Figure 2I**) and, as expected, SH had among the highest disome scores for viral mRNAs.

### RSV transcripts require eIF4E for translation

Our findings that viral mRNAs are efficiently translated and effectively compete with host mRNAs for ribosomes raises the question of whether they utilize any non-canonical mechanisms to recruit ribosomes. We therefore tested whether viral mRNAs require translation initiation factor eIF4E (**Figure S6A**) or utilize an alternative (non-canonical) mechanism to initiate translation. We performed ribosome profiling of eIF4E depleted cells during RSV infection (**Figure 3A**). We confirmed eIF4E knockdown by western blot (**Figure S6B**). When lacking phosphorylation, 4E binding proteins (4E-BPs) serve as a negative regulator of eIF4E by sequestering it and reducing overall translation (**Figure S6A**) (Ma and Blenis 2009). Drastic eIF4E knockdown has been shown to evoke a compensatory mechanism through degradation of 4E-BP (Yanagiya et al. 2012). We therefore checked 4E-BP1 levels after eIF4E knockdown to test whether this mechanism was counteracting the knockdown and observed only a mild decrease (**Figure S6C**). As more direct evidence of the knockdown’s effectiveness, we also tested the extent of translation inhibition by polysome profiling and confirmed a more than 2-fold decrease in polysome:monosome ratio (**Figure S6D**), indicating that the eIF4E knockdown was effective.

**Figure 3.**
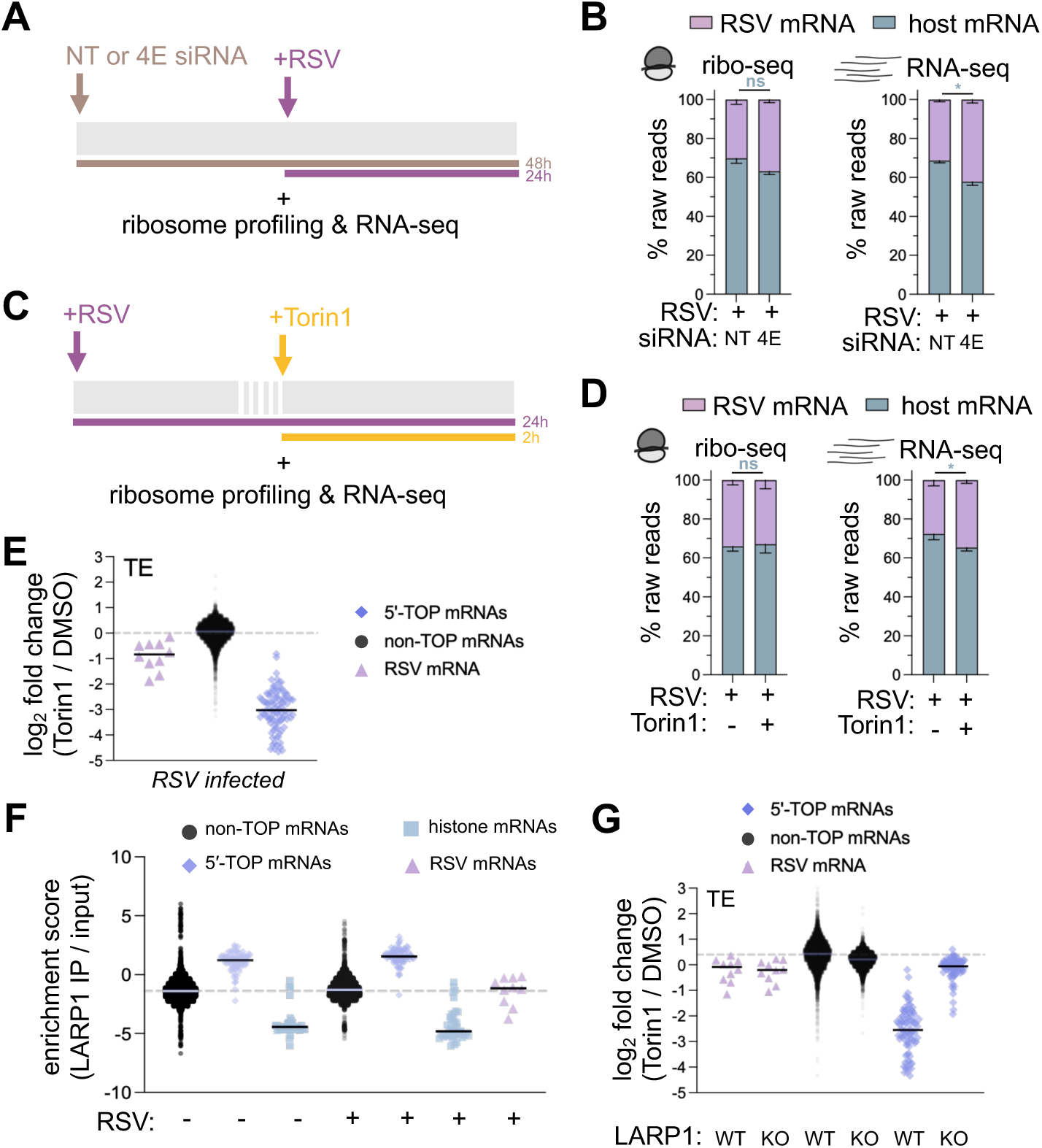
mTOR inhibition reduces the TE of RSV mRNAs independent of LARP1. (A,C) Schematic of experimental set-up. Transfection of eIF4E siRNA was done 24 hours before RSV infection (MOI 3) (A). Torin1 treatment was done in the final 2 hours of RSV infection (MOI 3) (C). (B,D) Quantification of the relative abundance of host and viral mRNAs for ribo-seq and RNA-seq datasets, shown as a percentage of raw read counts. Loss of eIF4E or treatment with Torin1 had a small effect that led to a slight relative increase in viral mRNAs (RNA-seq data). A paired t-test was performed to compare host mRNA abundance (blue bars) between eIF4E siRNA and non-targeting siRNA (NT) transfected cells (B) and Torin1 and DMSO treated cells (D). Bars represent the mean and error bars indicate the standard deviation. (E) Effect of Torin1 treatment in infected cells on TE computed by DESeq2. The TE of 5′-TOP mRNAs was strongly decreased, and viral mRNA TE showed a modest decrease. (F) LARP1 enrichment scores were calculated by dividing counts from LARP1 immunoprecipitation (IP) by input counts (in TPM units). Histone mRNAs were not enriched since these mRNAs lack a poly(A) tail, which is known to have some affinity for LARP1 and therefore serve as a negative control. RSV mRNAs cluster with non-TOP host mRNAs. (G) Differential TEs calculated for all individual mRNAs to determine the effect of Torin1 treatment during infection on WT and *LARP1* KO cells. The TE change of 5′-TOP mRNAs was dependent on LARP1. The small change in RSV mRNA TE due to Torin1 was not dependent on LARP1. P values < 0.05 are indicated by one asterisks (*) and ns indicates not significant.

To determine if RSV mRNAs depend on eIF4E for translation, we calculated the relative share of host and viral reads before and after loss of eIF4E in ribo-seq and mRNA-seq data (**Figure 3B**). We found that viral reads were slightly more abundant in both mRNA- and ribo-seq datasets, suggesting that RSV mRNAs may be somewhat more abundant upon depletion of eIF4E but lack any relative advantage in translation. We also measured viral protein levels by western blot and found they did not change much relative to GAPDH following eIF4E depletion (**Figure S6E**), consistent with the idea that RSV gene expression does not benefit from eIF4E depletion.

### mTOR inhibition reduces RSV mRNAs TE independently of LARP1

Regulation of eIF4E is under control of serine/threonine protein kinase mTORC1. Under normal growth conditions, mTORC1 is active which results in phosphorylation of 4E-BP1 and, as noted above, reduces its affinity for eIF4E (**Figure S7A**) (Gingras et al. 1999, 2001). A stress-inducing agent that deactivates the mTORC pathway is Torin1 (Thoreen et al. 2009). We tested the mTOR signaling pathway following two hours of Torin1 treatment and confirmed near complete dephosphorylation of 4E-BP1 by western blot (**Figure S7B,C**). Interestingly, consistent with a previous study (Pérez-Gil et al. 2015), we found that RSV also induced some 4E-BP1 dephosphorylation (**Figure S7B,C**). Next, to determine the effect of full mTORC deactivation on RSV infection, we harvested RSV-infected cells treated with Torin1 for 2 hours and performed ribosome profiling and RNA sequencing (**Figure 3C**). We found that the viral share of ribosome footprints was unchanged after Torin1 treatment (**Figure 3D**, ribo-seq). However, viral mRNAs became slightly more abundant (**Figure 3D**, RNA-seq, 28% to 35%). As a result, the translation efficiency (ratio of ribo-seq over RNA-seq calculated for each viral mRNA by DESeq2) decreased somewhat for viral mRNAs due to mTOR inhibition (**Figure 3E, Figure S7D, Table S7**). In particular, we found that the downregulation of 5 viral mRNAs met our significance threshold, including RSV-NS1, -NS2, -P, -N, and -G (**Figure S7E**, padj < 0.05 & FC < 1.5, **Table S7**). Torin1 treatment is also known to specifically inhibit the translation of 5′-TOP mRNAs, while also stabilizing these transcripts, through a LARP1-dependent pathway (Hochstoeger et al. 2024). We tested this expectation by examining the effects of Torin1 on 5′-TOP mRNAs and confirmed they were somewhat stabilized and their translation was decreased in our dataset (**Figure 3E, Figure S7D, Table S7**).

Regulation of 5′-TOP mRNAs occurs through specific binding by the RNA-binding protein LARP1 through their 5′-TOP motifs. We therefore employed several tests to determine whether the effect of Torin1 on the TE of viral mRNAs was also dependent on LARP1. First, we immunoprecipitated endogenous LARP1 and looked for enrichment of viral mRNAs. As a control, we confirmed specific enrichment of LARP1 by western blot (**Figure S8A**, compare LARP1 against IgG elution lanes). We then sequenced the RNA bound to LARP1 immunoprecipitation and calculated an enrichment score by dividing the LARP1-bound mRNA level by total mRNA level (in TPM, **Table S8**). Since LARP1 also binds poly(A) tails, we investigated histone mRNAs since they lack a poly(A) tail. As expected, these transcripts showed low association with LARP1 (**Figure 3F**). While the enrichment score for 5’-TOP mRNAs was high, as expected, we found that the scores for RSV mRNAs were similar those for non-TOP host mRNA (**Figure 3F**).

Next, we checked to see if the TE changes in viral mRNAs induced by Torin1 (see **Figure 3E**) were dependent on LARP1. We therefore generated CRISPR-Cas9 knockout cell lines and confirmed LARP1 KO by western blot (**Figure S8B**). As expected, the TE and abundance of 5’-TOP mRNAs was similar to non-TOP mRNAs in the absence of LARP1 (**Figure 3G, Figure S8C, Table S9**) (Philippe et al. 2020). In contrast, we found that the TE of viral mRNAs was not affected (**Figure 3G, Figure S8C**). Overall, these data show that during mTORC1 inhibition, RSV mRNAs are translated somewhat less efficiently and stabilized, and this effect is not dependent on LARP1. This is consistent with the absence of a 5’-TOP motif in at the start of the viral mRNAs (*i.e.* the transcription start site, TSS) (**Figure S8D**).

### RSV transcripts do not escape ISR-induced translational repression

Activation of the integrated stress response (ISR) can result in eIF2α phosphorylation through four kinases (**Figure 4A**), resulting in a global reduction in translation (see **Supplementary Figure 2A,B**). To determine if RSV transcripts are affected by ISR activation, we conducted ribo-seq and RNA-seq experiments on ISR-induced cells (through arsenite and thapsigargin treatment) (**Figure 4B**). We found ISR induction did not change the ratio of host and viral reads in either type of experiment, indicating that RSV mRNAs are as sensitive to ISR activation as are host mRNAs (**Figure 4C**). In addition, detection of nascent proteins following arsenite treatment of RSV infected cells eliminated protein production, consistent with the idea that ISR induction is not beneficial for RSV (see **Supplementary Figure 2B**). In contrast, poliovirus has been shown to maintain viral translation following arsenite treatment (White et al. 2011). Next, we measured eIF2α phosphorylation levels and, as a positive control, showed a strong increase in eIF2α-P following arsenite treatment. In addition, consistent with our previous study (Kerkhofs et al. 2025), we observed a very mild activation of eIF2α-P following RSV infection (**Figure 4D,E**), consistent with the model that its induction would adversely affect translation of the viral mRNAs.

**Figure 4.**
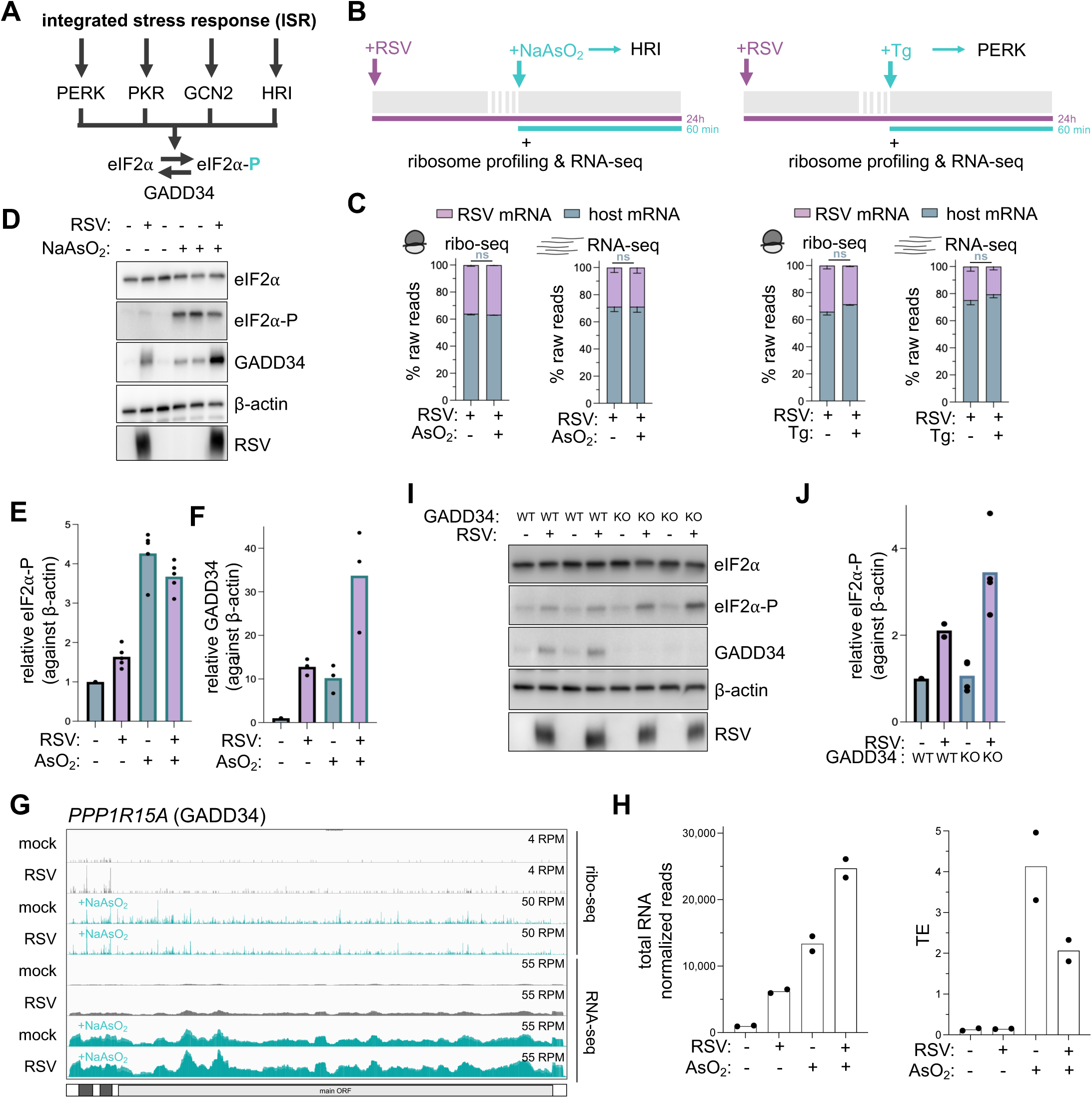
RSV transcripts are affected by ISR-mediated translational repression. (A) Schematic of kinases involved in eIF2α phosphorylation upon activation of the integrated stress response (ISR). Phosphorylation of eIF2α induces translational upregulation of the mRNA encoding the negative regulator GADD34 (also referred to as PPP1R15A). (B) Schematic of experimental set-up. Sodium arsenite (AsO_2_) and thapsigargin (Tg) treatment was done in the last hour of RSV infection (MOI 3). (C) Quantification of the relative abundance of host and viral mRNAs for ribo-seq and RNA-seq datasets, shown as a percentage of raw read counts. A paired t-test was performed to compare host mRNA abundance (blue bars) between ISR-inducing drug treatment and control treated conditions. Bars represent the mean and error bars indicate the standard deviation. (D-F) Western blot analysis detecting eIF2α phosphorylation and GADD34 levels (D). β-actin and total eIF2α served as loading controls and RSV protein was detected to confirm infection. RSV infection induces mild eIF2α phosphorylation and strong GADD34 upregulation, while arsenite treatment induced strong eIF2α phosphorylation and GADD34 upregulation, as quantified (E-F). Each datapoint indicates the value from a biological replicate. Combined arsenite treatment and RSV infection resulted in enhanced production of GADD34 protein, indicating activation through separate regulatory pathways. (G,H) IGV browser traces showing ribo-seq and RNA-seq data for the *PPP1R15A* (*GADD34*) mRNA (G). *GADD34* total mRNA levels were increased following both arsenite treatment and RSV infection and combined treatment showed an added effect, as quantified (H). The TE of *GADD34* mRNA only increased following arsenite treatment, as quantified (H). Each datapoint indicates the value from a biological replicate. Only arsenite treatment resulted in an increase in TE of *GADD34*. (I,J) Western blot analysis comparing eIF2α phosphorylation between wild-type (WT) and GADD34 knockout (KO) cells following RSV infection. β-actin and total eIF2α served as loading controls, GADD34 as knockout control and RSV protein was detected to confirm infection (I). GADD34 KO resulted in increased α phosphorylation, only during RSV infection, as quantified (J). Each datapoint indicates the value from a biological replicate. ns indicates not significant.

### GADD34 decreases eIF2α phosphorylation during infection

We next investigated the role of negative regulator GADD34 during infection (**Figure 4A**). We observed that both RSV infection and arsenite treatment strongly upregulated GADD34 protein levels (**Figure 4D,F**). We then used our ribosome profiling and RNA sequencing datasets to determine if GADD34 is upregulated at the mRNA or translational level during RSV infection. Arsenite treatment resulted in both transcriptional and translational (higher TE) upregulation (**Figure 4G,H, Table S10**). On the other hand, RSV infection induced GADD34 strictly transcriptionally (**Figure 4G,H**). Interestingly, these results appeared to be additive since combined arsenite treatment and RSV infection resulted in enhanced upregulation of GADD34 (**Figure 4D** and **F**; RSV + AsO2), likely reflecting the fact that GADD34 transcription is known to be activated by both the ISR and IFNs (Y. Ma and Hendershot 2003; Ruggieri et al. 2012; Lee et al. 2009).

Since stress-induced GADD34 upregulation results in eIF2αdephosphorylation, we tested the role of GADD34 in maintaining low eIF2α-P levels (and thus high translation levels) during viral infection. We therefore infected GADD34 KO cells with RSV and found that eIF2α-P levels were higher during RSV infection when GADD34 was absent (**Figure 4I,J**). We also tested whether individual domains of GADD34 played particular roles in RSV infection. As a positive control, we performed a rescue experiment and showed that eIF2α-P levels were reduced by overexpressing wild-type GADD34 in RSV infected KO cells (**Figure S9C**). Next, we performed the rescue experiment with deletion mutants of GADD34 to test its phosphatase activity. Upon deletion of the PP1 binding domain, we observed no reduction in eIF2α-P, indicating that the interaction between GADD34 and phosphatase PP1 is required for its function. Next, deletion of the PEST1 domain, which is required for interaction with eIF2α-P only partially rescued eIF2α-P levels. Since the ΔPEST1 domain GADD34 mutant retained some eIF2α binding activity (Choy et al. 2015), a portion of eIF2α-P can still be dephosphorylated. These findings indicate that GADD34 requires its canonical activity for reducing eIF2α-P during RSV infection. Overall, we found that GADD34 is transcriptionally upregulated during RSV infection and is important in maintaining low levels of eIF2α-P.

### RSV infection triggers the innate immune response without activating PKR and RNase L

Since RSV induces slight phosphorylation of eIF2α (see **Figures 4D, 4I and Figure S9C**), we next asked whether this was due to the action of PKR, which has been reported to be activated by dsRNA (Lemaire et al. 2008) and RSV infection (Lindquist et al. 2010b; Groskreutz et al. 2010). We determined PKR phosphorylation status through western blotting using a Phos-Tag gel. We transfected the double stranded RNA mimic poly(I:C) as a positive control for PKR activation and observed the expected gel shift (**Figure 5B**, upward shift caused by phosphorylation). In contrast, RSV infected cells did not induce PKR activation (**Figure 5B**). To exclude low level activation of PKR (below the detection limit of Phos-tag gel), we compared eIF2α phosphorylation levels between WT and PKR KO cells during infection. Interestingly, we found no difference between WT and KO cells (**Figure 5C**, compare lanes 3 and 4), suggesting other ISR inducing kinase(s) (Gcn2, PERK and/or HRI) are responsible for eIF2α phosphorylation during RSV infection (see **Figure 4A**).

**Figure 5.**
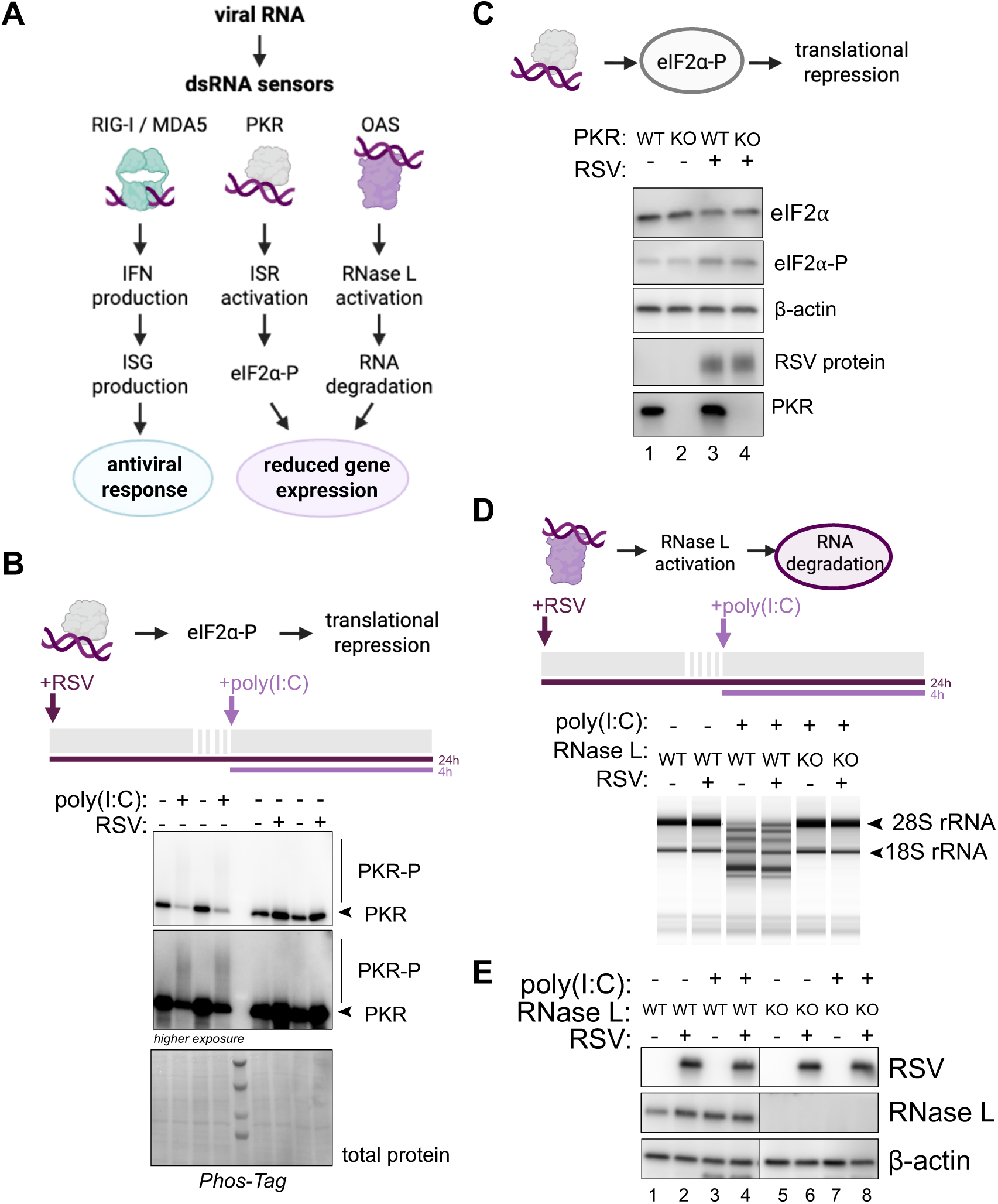
RSV infection activates IFN signaling without activating dsRNA sensors PKR and OAS. (A) Schematic representation of dsRNA sensors and downstream signaling pathways. RIG-I and MDA5 receptors induce IFN and ISG expression, leading to an antiviral response which is induced by RSV (see Figure S1). PKR activation leads to activation of the ISR and reduces global translation. Activation of OAS results activation of the endonuclease RNase L, leading to global RNA degradation. (B) Western blot analysis of Phos-Tag gel to assess PKR activation. PKR phosphorylation results in slower migration on Phos-Tag gels, allowing identification of activated PKR. PKR was detected by probing with a PKR antibody detecting total protein (and not PKR-P). Poly(I:C) transfection functioned as a positive control for PKR activation. Total protein was visualized by staining the membrane with Ponceau S staining. (C) Western blot analysis comparing eIF2αphosphorylation between wild-type (WT) and PKR knockout (KO) cells following RSV infection. Antibodies for β-actin and total eIF2α served as loading controls, antibody for PKR confirms KO, and antibody to RSV protein confirms infection. PKR deletion did not change eIF2α phosphorylation levels during RSV infection. (D) rRNA degradation detection by TapeStation analysis in WT and *RNASEL* KO cells. Poly(I:C) transfection functioned as a positive control for OAS activation, seen as an rRNA degradation pattern that disappeared in *RNASEL* KO cells. RSV infection did not induce OAS activation. (E) Western blot analysis of RNase L in WT and *RNASEL* KO cells confirmed absence of RNase L protein and upregulation under stress. RNase L is an ISG and is upregulated following RSV infection and poly(I:C) transfection. β-actin served as a loading control. RSV protein levels remain similar following RNase L removal.

To test if RSV infection specifically inhibits PKR activation or rather generically blocks dsRNA recognition, we tested activation of another cytoplasmic dsRNA receptor in our cells, OAS. Activation of OAS results in activation of the endonuclease RNase L which leads to global mRNA degradation (**Figure 5A**). RNase L activation results in specific cleavage within 18S and 28S rRNA which can be detected by an rRNA cleavage assay (Cooper et al. 2015; Karasik et al. 2021). We transfected poly(I:C) as a positive control for RNase L activation and observed the expected cleavage pattern, which disappeared in *RNASEL* KO cells (**Figure 5D**, compare lanes 1, 3 and 5) (Karasik et al. 2021). RSV infection did not result in rRNA cleavage (**Figure 5D**, compare lanes 1 and 2), consistent with previous work (Hwang et al. 2026). We also noted that poly(I:C) induced rRNA cleavage is not affected by RSV infection, indicating that that the virus does not employ any RNase L inhibition strategy (**Figure 5D**, compare lanes 3 and 4). In addition, to rule out low level of antiviral RNase L activation that specifically targets viral mRNA, we measured viral protein levels in the absence of RNase L. We observed the same amount of viral protein produced in WT and *RNASEL* KO cells, relative to β-actin, consistent with the observation that RNase L is not induced during RSV infection (**Figure 5E**, compare lanes 2 and 6). Note that RNase L is an ISG and its protein levels are increased during RSV infection and following poly(I:C) treatment (**Figure 5E**, compare lanes 1-4) (Panuska et al. 1995; Zhou et al. 1993; Pandey et al. 2004). While this indicates the IFN response is active (see **Figure S1D**) and upregulates *RNASEL*, we conclude the increased RNase L must not be active due to insufficient OAS activity. Overall, we demonstrate that RSV does not induce RNase L-induced RNA fragmentation. This strategy that would appear to be important since RSV mRNAs would be vulnerable to activation (Drappier and Michiels 2015).

In **Figure S1D**, we showed that RSV infection results in upregulation of IFNs and ISGs during RSV infection, consistent with prior work (Czerkies et al. 2022; Selvaggi et al. 2014). This is consistent with RSV activating innate immune receptors RIG-I and MDA5 (**Figure 5A**) (Liu et al. 2007; Grandvaux et al. 2014), without activating PKR and OAS. Since innate immune receptors RIG-I and MDA5 are not strictly activated by dsRNA, it is possible that an RSV-induced single stranded 5’-triphophosate RNA activates these receptors without activating OAS and PKR. This implies that the strategy adopted for evading PKR and OAS activation is not sufficient for avoiding other sensors that can detect dsRNA in the cell.

## Discussion

A longstanding challenge for understanding viral infection is the mechanism by which viruses gain access to the limited translational machinery and avoid activation of immune defense pathways. Many viruses induce host shutoff to specifically reduce host gene expression, dampening the immune response, and enhance viral access to the translational machinery (Rozman et al. 2023). However, it has remained unclear whether this sort of strategy is utilized by RSV. Resolving this question is important for more broadly understanding how RSV reprograms the host cell to facilitate its own propagation and, ultimately, developing better therapeutics that target the virus.

To address this challenge, we performed RNA-seq, with spike-in controls for normalization, to assay absolute RNA levels during viral infection. We also used ribosome profiling and an innovative spike-in based method for measuring RNA levels across a sucrose gradient to assay how the virus changed global translation. Our approach revealed many features of the RSV genetic program. Notably, RSV does not appear to employ any host shutoff mechanisms to limit the expression of host genes. In particular, we found that RSV does not take steps to reduce the level of host RNA in the cell, in contrast to VACV, where host mRNAs are strongly degraded (Ly et al. 2022). RSV manages to generate enough mRNA to constitute about 20% of the total transcriptome and slightly increases the overall level of RNA in the cell. We found that the translation of viral RNA is efficient, on par with that of well-translated host RNAs, which also led to ribosome collisions. Viral translation, which utilized about 30% of the ribosomes in the cell, was also steady over the course of infection. In contrast, the translation of SARS-CoV-2 mRNAs was reported to decrease as infection progressed (Finkel et al. 2021). The authors hypothesized that this is caused by the formation of double-membrane replication compartments that capture viral mRNAs, preventing them from being translated. While RSV also produces separate replication compartments (also known as inclusion bodies), these are formed early during infection and remain throughout infection (Ratnayake et al. 2026; Rincheval et al. 2017).

We also explored whether the virus used any unusual strategies to favor translation of its own RNAs, an approach utilized by other viruses, such as poliovirus (Kwan and Thompson 2019). In particular, we tested whether loss of inhibition of cap-binding protein eIF4E changed viral gene expression and found the virus depended on it, much like the host. Interestingly, we found that loss of eIF4E slightly increased viral RNA levels, relative to host, and inhibition of eIF4E through Torin1 slightly suppressed their translation. While these broad trends are reminiscent of the effects of the 5’-TOP host RNAs, we found the mechanism underlying this trend did not rely on the ribosome homeostasis sensor LARP1 (Saba et al. 2026), which is known to control these features of 5’-TOP mRNAs. Overall, we found viral mRNAs did not employ any obvious mechanism to circumvent the canonical mechanism of translation. Recent work in HAP1 cells (chronic myelogenous leukemia) showed that several negative-sense single-stranded RNA viruses, including RSV, use the alternative 5’-cap binding protein eIF4E3 to maintain viral translation when canonical eIF4E is depleted (Grimins et al. 2026). However, eIF4E3 levels are relatively low in A549 cells that were used in this study (Grimins et al. 2026). A549 cells (lung carcinoma cell) have eIF4E3 mRNA levels comparable to alveolar macrophages and explanted human lung tissue (Bertrams et al. 2022), suggesting that this effect may be cell type dependent. Last, we checked the role of host antiviral pathways ISR and OAS/RNase L interacted with the virus. We found the virus only minimally activated the ISR and did not activate OAS. However, it also did not block their activation by exogenous means (*i.e.* poly I:C transfection). Consistent with the view that the virus does not use non-canonical mechanisms to express it genes, these data suggest it is important that the virus avoid activation of these pathways, presumably by hiding its dsRNA.

RSV therefore employs a “fair competition” based approach in that its RNAs are abundant and efficiently translated and are sufficient to create viral proteins, despite host gene expression continuing in the background. Whether or not the ongoing host mRNA translation is important for the virus remains an open question. In addition, the ability of the virus to largely evade the ISR and completely evade OAS activation are critical consequences of this approach since the virus does not appear to have the ability to inhibit the pathways or avoid their consequences in terms of reduced gene expression.

Some of the phenotypes of eIF4E inhibition either via knockdown or treatment with Torin1 (and 4E-BP activation) suggest potential regulatory mechanisms for the viral RNA. One possibility is that turnover of the viral genes is somewhat dependent on translation, explaining why loss of translation initiation results in their stabilization. Many pathways could drive this phenomenon, such as nonsense mediated decay (NMD) or codon-optimized mediated decay (COMD, (Linder et al. 2025; Murakami et al. 2025; Thermann et al. 1998)). However, it is unlikely these pathways are responsible since RSV mRNAs lack obvious premature termination codons or exon-junction complexes (associated with NMD) and COMD has been shown to rely on m6A modification (Linder et al. 2025) but RSV mRNAs become less abundant when the m6A modification is reduced (Xue et al. 2019)

It is also intriguing to note that the ISG GADD34 is a negative regulator of the ISR and induced by interferon (and therefore presumably performs an antiviral function normally). As expected, we observed that GADD34 loss led to higher levels of eIF2α-P, which would be disadvantageous for the virus since the virus relies on the canonical translation machinery. It therefore remains an open question how GADD34 facilitates an anti-viral function against viruses like RSV that do not induce host shutoff.

Together, our data provide a full portrait of how RSV effectively competes with host mRNAs for the translational machinery to express its genes. Our study therefore offers an important window on the types of strategies that would be effective at therapeutically treating the virus.

## Methods

### Cell culture and generation of CRISPR-Cas9 modified cell lines

WT, *LARP1, PPP1R15A* (*GADD34*), *EIF2AK2* (*PKR), and RNASEL* A549 KO cells were maintained in RPMI 1640 GlutaMAX media (Gibco 61870127) supplemented with 10% FBS (Gibco A3160502). HEp-2 cells (ATCC, CCL-23) were maintained in DMEM media (Gibco 11965118) supplemented with 10% FBS. All cells were kept in a humidified incubator at 37°C with 5% CO_2_. All cell lines were routinely screened for *Mycoplasma* contamination and confirmed to be negative throughout the study using the Myco-X™ Mycoplasma PCR Detection Kit (BioLink). The WT, *EIF2AK2 and RNASEL* KO A549 cell lines were a kind gift of Dr. Bernie Moss and were generated as described (R. Liu and Moss 2016). The *GADD34* A549 KO (in an *RNASEL* KO background) and matching *RNASEL* A549 KO cell line was a kind gift of Dr. James Burke and was generated as described (Burke et al. 2020).

To generate the A549 *LARP1* KO cell line, we received the pX458 plasmid containing CRISPR-Cas9–targeting exon 5 of the LARP1 gene as a kind gift from Dr. Antonio Gentilella, as described (Fuentes et al. 2021). We prepared to transfect confluent WT A549 cells in 60-mm plates containing 3 mL media by adding 10 µg pX458 plasmid to 500 µL OptiMEM (Gibco 31985070) and adding 25 µL Lipofectamine 2000 Transfection Reagent (Invitrogen 11668027) to 500 µL OptiMEM. Following a 5-minute incubation at room temperature, the Lipofectamine-OptiMEM mixture was added to the plasmid-OptiMEM mixture and gently pipetted to mix. After 20-minute incubation at room temperature, the entire mixture was added dropwise to the cell media and incubated for 24 hours. Next, single-cell sorting of GFP-positive cells was performed at the NHLBI Flow Cytometry Core (NIH). Individual clones were grown in 96-well plates containing RPMI 1640 GlutaMAX media supplemented with 20% FBS and 1% Penicillin-Streptomycin (Gibco 15140122). Subsequently, the clones were propagated in RPMI supplemented with 10% FBS. Reduction of LARP1 expression in individual clones was tested by western blotting, resulting in identification of a LARP1 KO A549 cell line (clone #34).

### siRNA knockdown

All siRNAs were purchased from Horizon Discovery as ON-TARGETplus SMARTPools with the following catalogue numbers: eIF4E (L-003884-00-0005) and non-targeting (D-001810-10-05). The siRNA pools were resuspended to 100 µM in water and stored at - 80°C. We prepared to transfect confluent WT A549 cells in 100-mm plates containing 10 mL media by adding 1.5 µL 100 µM siRNA to 750 µL OptiMEM (Gibco 31985070) and 45 µL Lipofectamine RNAiMAX Transfection Reagent (Invitrogen 13778075) to 750 µL OptiMEM. Following a 5-minute incubation at room temperature, the RNAiMAX-OptiMEM mixture was added to the siRNA-OptiMEM mixture and gently pipetted to mix. After 20-minute incubation at room temperature, the entire mixture was added dropwise to the cell media and incubated for 6 hours. Next, cells were seeded in appropriate cell culture vessels for experiments. Cells were harvested 48 hours post-transfection and reduction of eIF4E expression was tested by western blotting **(Figure 3 and S6)**.

### Poly(I:C) and GADD34 plasmid transfection

We transfected confluent WT and *RNASEL* KO A549 cells in 6-well plates with 0.25 μg poly(I:C) per well (Invivogen tlrl-picw) for 4 hours. To prepare for transfection, poly(I:C) was resuspended in 125 µL OptiMEM (Gibco 31985070) and 10 µL Lipofectamine 2000 Transfection Reagent (Invitrogen 11668027) was added to 125 µL OptiMEM. Following a 5-minute incubation at room temperature, the Lipofectamine-OptiMEM mixture was added to the poly(I:C)-OptiMEM mixture and gently pipetted to mix. After 20-minute incubation at room temperature, the entire mixture was added dropwise to the cell media and incubated for 4 hours (**Figure 5**).

GADD34 expression plasmids were obtained from Addgene (WT: 75478, ΔPP1-dom: 75479, ΔPEST1: 78096) (Choy et al. 2015) and plasmid DNA was prepared using the QIAGEN Plasmid Mini Kit (Qiagen 12143). Plasmid DNA concentration was determined by Nanodrop spectrophotometer (Thermo Fisher) and adjusted to 1 µg/µL. For transfections of *GADD34* KO A549 cell lines in 12-well plates, 1 µg plasmid DNA was resuspended in 100 µL OptiMEM (Gibco 31985070) and 2.5 µL Lipofectamine 2000 Transfection Reagent (Invitrogen 11668027) in 100 µL OptiMEM. Following a 5-minute incubation at room temperature, the Lipofectamine-OptiMEM mixture was added to the plasmid-OptiMEM mixture and gently pipetted to mix. After 20-minute incubation at room temperature, the entire mixture was added dropwise to the cell media and incubated for 48 hours **(Figure S9)**. Protein samples for experiments were then prepared according to the western blot analysis protocol below.

### RSV infections

The RSV A2 P3 virus stocks were prepared by infecting 80% confluent HEp-2 cells with RSV A2 P2 (MOI 0.05) for 2 hours at 37°C. Following infection, the P2 inoculum was removed, and the cells were maintained in DMEM containing 2% FBS for about 3 days until syncytia formation. To harvest the P3 virus, the cells were scraped and collected with the cell culture media (containing released virus). The cell suspension was vortexed to release cell-associated virus and centrifuged for 15 minutes at 300g. The RSV stock P3 was aliquoted, snap frozen in liquid nitrogen and stored at −80°C.

RSV titrations were performed on confluent HEp-2 cells in 24-well plates as described previously (Luongo et al. 2013). In brief, we determined RSV titers by preparing 10-fold dilutions of the virus stock in DMEM in quadruplicate. Cells were infected for 2 hours followed by addition of 1 mL 0.8% methylcellulose in OptiMEM overlay. Following 7 days of infection, plates were fixed with 80% ice-cold methanol twice, blocked in 5% Blotting-Grade Blocker (Bio-Rad 1706404) in PBS and individual plaques were visualized by immunostaining with primary goat polyclonal anti-RSV antibody (Medix Biochemica V0601), secondary anti-goat HRP-coupled antibody (BioRad 1721034) and KPL TrueBlue Peroxidase Substrate (SeraCare 5510-0030).

Experiments were performed with the RSV A2 strain (ATCC, serial passage-3 (P3)) and added to cells at an MOI 3 and this generally achieved an infection >90%. The aliquoted virus was thawed quickly at 37°C and diluted in FBS-free RPMI 1640 GlutaMAX media to an MOI of 3. Prior to infection, the monolayer was washed once with PBS, followed by incubation with a small volume of FBS-free inoculum (i.e. 15 cm plates: 5 mL, 10 cm plates: 2 mL, 24-well plates: 200 μL, 96-well plate: 32 μL). Infections were done for 2 hours at 37°C with frequent rocking to redistribute the inoculum evenly across the monolayer. Mock infections included PBS wash and 2 h incubation in FBS-free RPMI1640 GlutaMAX media. Following infection, cells were maintained in RPMI 1640 GlutaMAX media containing 10% inactivated FBS (30 minutes at 56°C) for 24 hours (unless stated otherwise).

### Drug treatments

Thapsigargin (Ambeed A594348) and Torin1 (Tocris Bioscience 4247) were dissolved in DMSO. Paired negative control cells were treated with an equal volume of DMSO. Cells were incubated with 250 nM Torin1 for 2 h or with 25 nM Torin1 for 24 h, 1 μM Thapsigargin for 1 h and 0.5 mM NaAsO_2_ (Millipore Sigma 1062771000) for 1 h. For serum starvation, the cells were washed with PBS, followed by incubation in RPMI 1640 GlutaMAX media without FBS supplement for 24 h.

### SUrface SEnsing of Translation (SUnSET) assay

For measuring nascent protein synthesis, cells were treated with 10 µg/mL puromycin (Sigma P9620) for 10 minutes, immediately followed by lysis following the western blot method below **(Figure S2)**. The negative control experiment testing inhibition of global protein synthesis was done by adding 100 µg/mL cycloheximide (Sigma C7698) to the media 10 minutes prior to puromycin treatment. Similarly, 0.5 mM sodium arsenite was added to the cells 50 minutes prior to puromycin treatment.

### LARP1 immunoprecipitation

Cells were grown to ∼70-80% confluency in 150-mm plates and washed with PBS prior to harvest. Ribonucleoprotein immunoprecipitation of LARP1-associated RNA was performed by in-plate crosslinking with 0.1% Formaldehyde in PBS (Thermo Scientific 28908) for 15 minutes at room temperature, followed by immediate in-plate quenching with 0.25 M glycine in PBS for 5 minutes. The cells were lysed by addition of 1 mL complete IP lysis buffer [50 mM HEPES, pH 7.5, 400 mM NaCl, 1 mM EDTA, 1 mM DTT, 0.5% Triton X100, 10% glycerol, 2 mM PMSF, 1X PIC, 200U/mL RNase inhibitor, 50 U/mL TurboDNase] and cell scraping. Lysates were collected in Eppendorf tubes and centrifuges for 10 minutes at 20,000 g at 4°C. The supernatants were collected and flash frozen in liquid N_2_ and stored at -80°C until use.

Lysates were thawed on ice, and 100 µL lysate was retained as the RNA input sample. Protein G Dynabeads (Invitrogen 10003D, 50 µL per condition) were coupled to 1 µg rabbit anti-LARP1 antibody (Abcam, ab86359) and 1 µg rabbit IgG control antibody (Cell Signaling Technology 2729) as a negative control, according to manufacturer’s instructions. Following bead washing to remove unbound antibody, 250 µL lysate was added and incubated for 2 h at 4°C on a nutator. The beads were then washed four times with wash buffer [50 mM HEPES, pH 7.5, 400 mM NaCl, 1 mM EDTA, 1 mM DTT, 0.5% Triton X100]. The final bead suspension was transferred to a new Eppendorf tube, followed by resuspension of the beads with 120 µL 1X RIP buffer [2X: 100 mM HEPES, pH 7.5, 200 mM NaCl, 10 mM EDTA, 20 mM DTT, 1% Triton X100, 20% glycerol, 2% SDS]. An equal volume of 2X RIP buffer was added to the input sample. Next, IP and input samples were incubated at 70°C for 45 min to reverse cross-linking, followed by RNA extraction using TriPure Isolation Reagent (Roche 11667165001), according to manufacturer’s instructions **(Figures 3 and S8)**.

### Sucrose gradient fractionation

Cells were grown to ∼70-80% confluency in 150-mm plates before harvest and sucrose gradient fractionation was performed as described in (Morita et al. 2013). Prior to collection, 100 μg/mL cycloheximide was added for 5 min at 37 °C. The cells were washed with PBS containing 100 μg/mL cycloheximide followed by trypsinization of the cells. Pellets were collected by centrifugation (5 minutes at 300g) following addition of RPMI 1640 GlutaMAX media containing 10% FBS and 100 μg/mL cycloheximide to neutralize trypsin. The cell pellet was washed twice with PBS and stored at −80°C until use.

Cell pellets were lysed in 485 μL hypotonic buffer [5 mM Tris–HCl (pH 7.5), 2.5 mM MgCl_2_, 1.5 mM KCl, 1X Halt™ Protease Inhibitor Cocktail, 100 μg/mL cycloheximide, 2 mM DTT, 200 Units/mL SUPERase In™ RNase Inhibitor, 0.5% (v/v) Triton X-100 and 0.5% (w/v) sodium deoxycholate, 85 μg heparin], followed by centrifugation for 5 min at 20,000 g at 4°C. Equal amounts of each sample (500 μL of A260=10) was loaded on a 7-step (sequentially frozen layers) 20–50% sucrose gradient prepared in sucrose buffer [20 mM HEPES (pH 7.6), 100 mM KCl, 5 mM MgCl_2_ and 100 μg/mL cycloheximide] in polypropylene centrifuge tubes (Beckman Coulter 331372, 14 × 89 mm). Next, the samples were ultracentrifuged (Beckman L-90K or XE-90) at 30,000 RPM for 3 hours in a Beckman SW41Ti rotor at 4°C with maximum acceleration and slow deceleration. The gradients were fractionated and collected (27 sec) using a Brandel Density Gradient Fractionation System, which monitored the RNA at 254 nm and digitally recorded polysome traces with a DataQ Instruments DI-155. Fractions were stored at −80°C or used immediately for RNA precipitation (**Figures 1-2, S4-S6)**.

Fractions were identified as 80S monosome, light polysomes (2-3 ribosomes), middle polysomes (4-5 ribosomes) and heavy polysomes (6+ ribosomes) based on the polysome traces. The bottom fraction of the gradient was not included in the heavy polysome fraction. ERCC spike-in containing TriPure Isolation Reagent (Roche 11667165001) was prepared by adding 1:100 diluted spike-in RNA to a concentration of 3.33 μL/mL, followed by vortexing. Each fraction contained 600 μL to which 600 μL spike-in TriPure Isolation Reagent was added. At this point, fractions were pooled and RNA was extracted according to the manufacturer’s instructions. rRNA depletion and library preparation was performed by the NHLBI DNA Sequencing and Genomics Core. Sequencing was performed on an Illumina Novaseq platform using paired-end 50 (PE50) sequencing at the NHLBI DNA Sequencing and Genomics Core.

### Library preparation for ribosome and disome profiling

Cells were treated and/or infected and harvested at 70-90% confluency in T75 flasks or 150-mm plates by washing the monolayers with ice-cold PBS and flash freezing the cell culture vessels directly in liquid N_2_ and stored at -80°C until use. The frozen cells were lysed by adding 600 or 1200 μL (T75 and 150-mm, respectively) ice-cold lysis buffer [20 mM Tris–HCl pH 7.4, 150 mM NaCl, 5 mm MgCl2, 1 mM DTT, 100 μg/ml cycloheximide, 1% Triton X-100, 25 U/mL Turbo DNase] while being thawed on ice. Cells were scraped and collected in Eppendorf tubes, followed by 10 minutes incubation on ice before being passed through 25G needles 10 times. The cell lysates were centrifuged for 10 minutes at 20,000 g at 4°C, flash frozen in liquid N_2_ and stored at -80°C until use. Library preparation of the lysates were performed as described before (McGlincy and Ingolia 2017). In brief, 300 μL lysate was incubated with 7.5 μL RNase I (100 U / μL, Invitrogen AM2294) for 1 hour at room temperature at 700 RPM. To halt the RNase digest, 10 μL of SUPERase·In RNase Inhibitor (20 U/μL, Invitrogen AM2696) was added and lysates clarified for 5 minutes at 20,000 g at 4°C. These lysates were ultracentrifuged on a sucrose cushion [1 M sucrose, 20 mM Tris-HCl pH 7.4, 150 mM NaCl, 5 mM MgCl_2_, 1 mM DTT, 100 μg/ml cycloheximide, 20U/mL SUPERase·In RNase Inhibitor] using a TLA100.3 rotor (100,000 RPM for 1h at 4°C) to pellet the ribosomes. The pellet was resuspended in 700 μL TriPure Isolation Reagent (Roche 11667165001) and extracted according to the manufacturer’s instructions. The RNA was quantified using the Nanodrop spectrophotometer (Thermo Fisher) and up to 10 μg RNA was loaded on a 15% Criterion TBE-urea polyacrylamide gel (BioRad 3450091) by mixing equal amount of sample with 2X loading dye [7 M urea, 0.12 g/mL Ficoll-400, 1X TBE]. The gel was cut to extract ribosome footprints sizes to include the 25-34 nucleotide range. RNA elution buffer [0.3 M sodium acetate, 1 mM EDTA, 0.25% SDS] was added to the gel slice, incubated for 30 minutes on dry ice, followed by elution of the RNA from the gel slice overnight at room temperature at 700 RPM. The RNA was precipitated by adding GlycoBlue™ Coprecipitant (15 mg/mL, Invitrogen AM9516) and a volume of isopropanol equal to the sample volume and incubated for 1 hour on ice. Next, the RNA was dephosphorylated for 1 hour at 37°C using 5U T4 PNK (10 U/μL, NEB M0201) and pre-adenylated linkers containing unique internal barcodes were ligated for 3 hours at 22°C using T4 RNA Ligate2 Truncated (K227Q) (NEB M0351) as described in (McGlincy and Ingolia 2017), followed by 5’-deadenylase (NEB M0331) and RecJ exonuclease (Lucigen RJ411250) treatment for 45 minutes at 30°C. At this point, samples with unique internal barcodes were pooled and rRNA depleted using the riboPOOL rRNA depletion kit (siTOOLS dp-K024-50).

Following rRNA removal, the RNA was reverse transcribed with SuperScript III Reverse Transcriptase (Invitrogen 18080044), as described in (McGlincy and Ingolia 2017) and loaded on a 10% Criterion TBE-urea polyacrylamide gel (BioRad 3450089) by mixing equal amount of sample with 2X loading dye [7 M urea, 0.12 g/mL Ficoll-400, 1X TBE] for sample purification. DNA elution buffer [0.3 M NaCl, 1 mM EDTA, 10 mM Tris-HCl, pH 8.0] was added to the gel slice, incubated for 30 minutes on dry ice, followed by elution of the RNA from the gel slice overnight at room temperature at 700 RPM. The DNA was precipitated by adding GlycoBlue™ Coprecipitant (15 mg/mL, Invitrogen AM9516) and a volume of isopropanol equal to the sample volume and incubated for 1 hour on ice. The cDNA was circularized using CircLigase ssDNA Ligase (Biosearch Technologies, CL4111K) according to manufacturer’s instructions. Finally, libraries were generated by PCR using Illumina barcode containing primers, as described in (McGlincy and Ingolia 2017). Quality of the libraries were assessed by TapeStation 4150 using DNA ScreenTape Analysis (Agilent 5067-5584). Sequencing was performed on an Illumina Novaseq platform using single-end 100 (SE100) sequencing at the NHLBI DNA Sequencing and Genomics Core **Figures 2-4, S3, S7-S8**.

Preparation of disome profiling samples were prepared according to the ribosome profiling protocol with following changes. RNase I digest was milder and performed by adding 2 μL RNase I (10 U / μL, 1:10 dilution of 100 U / μL stock, Invitrogen AM2294) per 10 μg RNA for 1 hour at room temperature at 700 RPM. During size selection on the 15% Criterion TBE-urea polyacrylamide gel (BioRad 3450091), the gel was cut to extract disome footprints (50-70 nucleotide range) and matching monosomes (25-34 nucleotide range) (**Figure 2**).

### Library preparation for RNA-seq (matching ribo-seq)

Total RNA was extracted from 100 μL ribosome profiling lysates using TriPure Isolation Reagent (Roche 11667165001) (according to the manufacturer’s instructions; cells grown in a monolayer option A). The extracted RNA was quantified using the Nanodrop spectrophotometer (Thermo Fisher) and RNA quality was evaluated by running the RNA on a TapeStation 4150 using RNA ScreenTape Analysis (Agilent 5067-5576), prior to library preparation.

For the RSV timecourse ERCC spike-in containing RNA-seq, the RNA concentration was measured by Nanodrop, diluted to 1.5 μg for each sample and measured again. To 1.5 μg of total RNA, 3 μL of a 1:100 diluted ERCC RNA Spike-In Mix (Invitrogen 4456740) was added. Next, the samples were rRNA depleted using the riboPOOL rRNA depletion kit (siTOOLS dp-K024-53), followed by library preparation using the NEBNext Ultra™ II Directional RNA Library Prep Kit for Illumina (NEB, E7760) with Illumina linker ligation using NEBNext Multiplex Oligos for Illumina (Index Primers Set 1, NEB E7335S), according to manufacturer’s instructions.

The remaining datasets (4E siRNA, LARP1 immunoprecipitation, Torin1 and arsenite treated) were prepared without ERCC spike-in addition, and rRNA depletion and library preparation was performed by the NHLBI DNA Sequencing and Genomics Core. The thapsigargin treated and LARP1 KO datasets were rRNA depleted using the NEBNext rRNA Depletion Kit v2 (NEB E7400) and libraries were prepared using the NEBNext UltraExpress RNA Library Prep Kit (NEB E3330), according to manufacturer’s instructions. Sequencing was performed on an Illumina Novaseq platform using paired-end 50 (PE50) sequencing at the NHLBI DNA Sequencing and Genomics Core.

### Read processing and alignment

#### General overview

The NHLBI DNA Sequencing and Genomics Core performed demultiplexing of Illumina barcodes, producing fastq files. Internal barcodes for ribo-seq datasets were demultiplexed using cutadapt. These were further processed and analyzed as detailed below. Fastq files from libraries run across multiple lanes/chips were combined using the *cat* command. All alignments were done with bowtie1 (Langmead et al. 2009). FastQC and MultiQC were used to perform quality controls throughout the processing and alignment steps. All read processing and alignment steps were performed on the NIH Biowulf cluster. Visualization in IGV was done by preparing bedgraph files for ribo-seq data and bigwig files for RNA-se data following Github instructions associated with the ribofootPrinter package (Kerkhofs and Guydosh 2026). To generate plots of ribo-seq 5’ ends or RNA-seq coverage, we used bedtools, deeptools, and IGV (Figure 4).

Additional details of code packages used and availability of custom python code on Github are detailed elsewhere.

#### Ribosome and disome profiling

The single-end fastq files were separated by unique internal barcodes and trimmed using Cutadapt (ribo-seq settings: *--discard-untrimmed --no-indels -e 1 -m 32 -M 41*, disome-seq settings: *--discard-untrimmed --no-indels -e 1 -m 57 -M 87*) (Martin 2011). Contaminating rRNA and tRNA sequences were removed by bowtie1 alignment (settings: *-v 2 -y -S --trim5 2 --trim3 5*) against the noncoding fasta file produced by downloading rRNA sequences from the SILVA project (release 128) (Quast et al. 2013) and tRNA sequences from GtRNAdb (*H. sapiens* release 16) (Chan and Lowe 2009). The fastq files containing unmapped reads were run through a custom Python script (dedup.py) to remove PCR duplicates by comparing the 7-nt unique molecule identifiers (UMIs) included in our libraries. Afterwards, the UMIs were trimmed using Cutadapt (settings: *-u 2 -u -5*).

Reads were aligned against the human MANE v1.4 transcriptome, as prepared in (Kerkhofs and Guydosh 2026), using bowtie1 (settings: *-v 1 -y --best -S -m 1*). Next, the unmapped reads were aligned against the RSV transcriptome (GenBank: KT992094.1) (**Additional Data File 1**) using bowtie1 (settings: *-v 2 -y --best -S -m 1*). The resulting SAM files were converted to BAM files and raw read counts were obtained using featureCounts (settings: *-t CDS -g gene_id -s 0 -M –fraction*) with gtf files for the human transcriptome from (Kerkhofs and Guydosh 2026) and the RSV transcriptome (**Additional Data File 2**). (**Table S11**). Fractional read counts were rounded to whole numbers for use with DESeq2 (see below).

#### RNA-seq

The paired-end fastq files were aligned against the human MANE v1.4 transcriptome using bowtie1 (settings: *-v 1 -y --best -S -m 1 -1 R1.fastq -2 R2.fastq*) and unmapped reads were aligned against the RSV transcriptome (GenBank: KT992094.1) (**Additional Data File 1**) using bowtie1 (settings: *-v 2 -y --best -S -m 1 -1 R1.fastq -2 R2.fastq*). For datasets containing ERCC spike-in RNAs, reads that failed to align against the host or viral transcriptome were then mapped to the ERCC transcriptome using bowtie1 without allow any mismatches (settings: *-v 0 -y -m 1 --best -S -1 R1.fastq -2 R2.fastq*). Raw read counts for spike-in mRNAs correlated (Pearson r) strongly with the known input concentrations of the individual spike-in mRNAs (r > 0.942) and between samples (r > 0.996), indicating the validity of the ERCC spike-in usage for subsequent normalization calculations.

The resulting SAM files were converted to BAM files and raw read counts were obtained using featureCounts (settings: *-t CDS -g gene_id -p -s 2 -M –fraction*) (**Table S11**) with gtf files for the human transcriptome from (Kerkhofs and Guydosh 2026) and the RSV transcriptome (**Additional Data File 2**). The VACV datasets generated in (Ly et al. 2022) were downloaded from SRP340405 on SRA and aligned against the human MANE v1.4, VACV and ERCC spike-in transcriptome following the same steps as described above. Fractional read counts were rounded to whole numbers for use with DESeq2 (see below).

### High-throughput sequencing data analysis

#### ERCC spike-in normalization

For the ERCC spike-in containing RSV timecourse (6h, 12h, 18h and 24h) and VACV timecourse (3h, 6h and 18h) datasets, raw read counts summed separately for host mRNAs, viral mRNAs and ERCC mRNAs. ERCC spike-in normalized read counts were then calculated by dividing the total read counts for host and viral mRNAs by the total read count for ERCC mRNAs. For data analysis without ERCC spike-in normalization, the percentage of raw reads corresponding to host and viral mRNAs was calculated by dividing the raw read count for each by the sum of host and viral mRNAs. The percentage of raw reads corresponding to ERCC spike-in mRNAs as calculated by dividing the raw read count for each by the sum of host, viral and ERCC mRNAs. Data was plotted in GraphPad Prism v10.6.0.

For datasets containing ERCC spike-ins from sucrose gradient, raw read counts were first converted to reads per kilobase (RPK) by dividing the raw read counts by the length of the main open-reading frame (ORF), in kilobases. Since longer mRNAs are expected to run deeper in the gradient, this normalization step accounts for longer mRNAs contributing more reads, enabling a more accurate comparison of ribosome loading. ERCC spike-in RNA was added proportionally to each fraction. However, the 80S monosome fraction consisted of one fraction, while light and middle polysomes consisted of two pooled fractions and heavy polysomes of three pooled fractions. To account for these differences in the number of pooled fractions, a weighting factor was incorporated by dividing the sum of the ERCC spike-in mRNAs by the corresponding number of fractions (80S: 1x, Lpol: 2x, Mpol: 2x, Hpol: 3x). In this way, the measurement is dependent on the species of ribosome in the fraction, and not impacted by how large a fraction is or how the concentration of sucrose affects RNA extraction efficiency. Afterwards, ERCC spike-in normalized read counts were calculated similarly as above. Data was plotted in GraphPad Prism v10.6.0.

#### mRNA distribution calculations, ribosome density and heatmap

To determine the distribution of each transcript along the polysome gradient, the percentage mRNA found in each of sucrose gradient fractions (80S, Lpol, Mpol and Hpol) was calculated. Similarly, the sum of the ERCC spike-in mRNAs was divided by the weighing factor to account for differences in the number of pooled fractions (see above) prior to ERCC spike-in normalization. Each host and viral transcript was then normalized by dividing its raw read count by the weighted sum of the ERCC mRNAs. Next, the ERCC spike-in normalized read counts were summed for each transcript across all four fractions, followed by dividing these reads counts by this sum to calculate the percentage mRNA distribution, such that the sum of each transcript adds up to 100%.

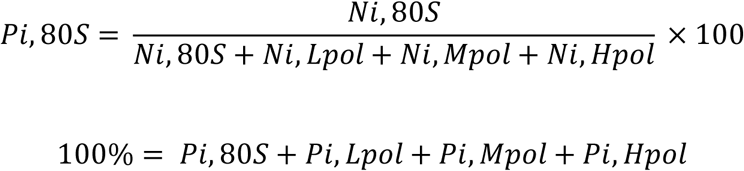

*Ni* = ERCC-normalized read count for transcript in fraction

*Pi* = percentage of transcript i in fraction

The mRNA distribution plots and matching pie charts for each individual viral mRNA, along with host mRNAs with similar ORF lengths (±30 nt) for comparison, were generated using a custom R script (*mRNA_distribution.Rmd*) and plotted using the R package *ggplot2 v4.0.1*. The percentage mRNA distribution data (as calculated above) was then used to determine the peak fraction in which each host mRNA was found most abundant using the *XLOOKUP* and *MAX* functions in Excel. The options for peak fraction include the 80S monosome, light, middle and heavy polysomes. The R input file containing percentage mRNA distribution data for each sample (as calculated above) and peak information was processed to generate plots for abundant transcripts (average ERCC normalized sum across fractions > 10) within the given ORF length range.

To generate the heatmap (**Figure S4**), transcripts were classified by main ORF length as short (< 500 nt), intermediate (between 500 and 1500 nt) and long (> 1500 nt). The R input file containing percentage mRNA distribution data for each sample (as calculated above) and ORF length information (short, mid, long) for each transcript was processed with a custom R script (*heatmap.Rmd*) to generate a heatmap of abundant transcripts (basemean > 100) using the *pheatmap v1.0.13* package in R.

To calculate ribosome density (note this is a fraction of all ribosome bound RNAs and does not include unbound RNAs), the percentage mRNA distribution data (as calculated above) was multiplied by a factor representing the average number of ribosomes associated with transcripts in each sucrose gradient fraction. The 80S monosome fraction was assigned a factor of 1, light polysomes (2–3 ribosomes) a factor of 2.5, middle polysomes (4–5 ribosomes) a factor of 4.5, and heavy polysomes (>6 ribosomes) an estimated factor of 10.5. The sum of these values across the gradient fractions represents the estimated average number of ribosomes associated with each transcript (ribosome count). The ribosome count was then divided by the corresponding ORF length to calculate ribosome density, represented as the number of ribosomes per 100 nt.

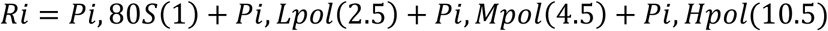

*Pi* = percentage of transcript *i* in fraction

*Ri* = the estimate average number of ribosomes for one molecule of transcript *i*

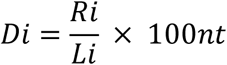

*Di* = the average ribosome density for transcript *i*

*Li* = the main ORF length for transcript *i*

#### Differential expression, TE and PCA analysis

Differential expression (DE) analysis for timecourse datasets (i.e. RNA-seq) was performed with the R package *DESeq2 v1.46.0* using raw read counts as inputs*. DESeqDataSetFromMatrix* was used to make comparisons for each timepoint separately between uninfected and infected conditions (design = ∼virus). For the ERCC spike-in containing RSV timecourse (6h, 12h, 18h and 24h, **Table S1**) and VACV timecourse (3h, 6h and 18h, **Table S2**) datasets, the ERCC spike-in raw read counts were used for normalization by applying *estimateSizeFactors* with *controlGenes*=ERCC. Volcano plots generated for RSV and VACV timecourse (**Figure S1D**) used padj and log2FoldChange calculated by DESeq2 including the ERCC normalization step and were made with the R package *ggplot2 v4.0.1*.

Translation efficiency (TE) from ribo-seq and RNA-seq experiments for the RSV infection timecourse dataset (**Figure 2C and Table S3**) was calculated using the R package *DESeq2 v1.46.0* using raw read counts as inputs. The RNA-seq and ribo-seq datasets for each condition (*e.g.* infected, 24h) were selected and comparisons to compute TE (ribo-seq/RNA-seq) were made using *DESeqDataSetFromMatrix* (design = ∼RNA).

Differential gene expression analysis for WT A549 Torin1 treated RNA-seq and ribo-seq datasets was performed using *DESeq2*, similar as above (**Table S7**, **Figure S7D**). *DESeqDataSetFromMatrix* was used to make comparisons between DMSO and Torin1-treatment for RSV-infected samples for (design = ∼treatment). Differential translation efficiency analysis was completed through setting the *DESeqDataSetFromMatrix* settings to obtain the ratio of Torin1 TE relative to DMSO TE (design=∼RNA + treatment + RNA:treatment). The plots comparing Torin1 vs DMSO TE in infected cells used padj and log2FoldChange calculated by DESeq2 and was generated with the R package *ggplot2 v4.0.1* (**Table S7, Figures 3E and S7E**).

Translation efficiency fold changes for datasets obtained for Torin1-treated RNA-seq and ribo-seq *LARP1* KO A549 cells were calculated using RPM normalized read counts and highly expressed transcripts were plotted (basemean > 10) (**Table S9 and Figures 3G and S8C**)

PCA analysis for the RNA-seq and ribosome profiling RSV timecourse (6h, 12h, 18h and 24h) datasets was performed by calculating the principal components by variance-stabilizing transformation (VST) of the DESeq2 raw count data (without ERCC spike-in normalization). Sample variance was visualized using the R package *ggplot2 v4.0.1* with PC2 plotted against PC1 (**Figure S3B**).

#### Ribosome profiling footprint size analysis

Analysis of mapped host and RSV mRNAs was performed using the Python toolbox ribofootPrinter2 using the *region_size_and_abundance.py* script (Kerkhofs and Guydosh 2026). This script uses the alignment SAM files and obtains information on the abundance and length of the mapped reads within a certain region within the transcript. The probability density for main ORF, start and stop codon was calculated for host mRNAs and RSV mRNAs to normalize each feature independently so that the total adds up to 1 (**Figure S3A**). This allows visualization of the read length distribution (without abundance) for each feature separately. The output Excel files for each sample of the RSV timecourse were parsed with a custom R script (generated by ChatGPT; *rsa_parser.Rmd*) to create a single file. The probability density data was extracted for main ORF, start and stop codon and the distribution graph was generated with a custom R script (generated by ChatGPT; *rsa_plot.Rmd*).

### Western blot analysis

Protein lysates were prepared by in-plate lysis of the cells. For this, the monolayers were rinsed once with PBS and an appropriate volume RIPA lysis buffer (Thermo Scientific 89900) containing 1X PIC (Thermo Scientific 78429) was added to the plates (*i.e.* 15 µL/cm^2^). Before lysates were collected, the plates were placed on ice for 20 minutes. Next, the lysates were centrifuged for 20 minutes at 20,000 g at 4°C.

Protein lysates were prepared for gel electrophoresis by incubation with 1× Laemmli buffer [5X Laemmli buffer: 5% β-mercaptoethanol (v/v), 0.02% bromophenol blue (w/v), 30% glycerol (v/v), 10% sodium dodecyl sulfate (SDS) (w/v), 250 mM Tris–HCl, pH 6.8] for 10 min at 95 °C. The samples were separated using a 4–20% gradient Mini-PROTEAN Tris–HCl gel (BioRad 4561096) for 35 min at 200V and transferred to a 0.2 μm PVDF membrane using the Trans-Blot Turbo System (BioRad, 1.3A, 7 minutes). Separation of proteins on a PhosTag gel, prepared according to manufacturer’s instructions [7% acrylamide/bis solution (29:1), 10.7 μM Phos-tag (Wako, AAL-107) and 21.3 μM MnCl_2_], was done for 1 hour at 150V. Prior to transfer, the Phos-Tag gel was washed 3 times in 10 mM EDTA, 10% methanol, and 1X Tris/Glycine transfer buffer (BioRad 1610734) and once in 10% methanol, and 1X Tris/Glycine transfer buffer. The proteins were then transferred to a 0.2 μm PVDF membrane using the Trans-Blot Turbo System (BioRad, 1.3A, 14 minutes). Membranes were blocked in 5% Blotting-Grade Blocker (Bio-Rad 1706404) in TBST (Tris-Buffered Saline with 0.1% Tween-20) for 1 hour or overnight at room temperature.

Next, the membranes were incubated in 1:2000 diluted primary antibody for 1 hour or overnight. Primary antibodies used in the study include goat anti-RSV (Medix Biochemica V0601), rabbit anti-GAPDH (Abcam ab9485), mouse anti-puromycin (Milipore Sigma MABE343), mouse anti-β-actin (Abcam ab8224), rabbit anti-eIF2α (Bethyl A300-721A), rabbit anti-eIF2α S51-P (Abcam ab32157), rabbit anti-eIF4E (Cell Signaling 9742), rabbit anti-4EBP (Cell Signaling 9644), rabbit anti-4EBP T37/T46-P (Cell Signaling 2855), rabbit anti-4EBP S65-P (Cell Signaling 9451), rabbit anti-LARP1 (Abcam ab86359), rabbit anti-GADD34 (Cell Signaling 41222), and rabbit anti-PKR (Abcam ab32052). After washing 5 times for 5 minutes in TBST, the membranes were incubated with 1:5000 diluted HRP-coupled anti-rabbit, anti-mouse or anti-goat secondary antibodies (BioRad 1706515, 1706516, 1721034) and incubated for 1 hour at room temperature. Following 5 washes in TBST, the PVDF membranes were incubated with Clarity Western ECL substrate (BioRad 1705061) for 5 minutes and proteins were visualized by Amersham Imager 600.

### Indirect immunofluorescent staining

Cells were washed with PBS, followed by fixation with 4% paraformaldehyde for 20 minutes and permeabilization for 10 min with 0.1% Triton X-100 in PBS. Cells were blocked for 1 hour in 1% Bovine Serum Albumin (BSA) in PBS, followed by incubation with primary antibodies in blocking solution for 1 hour at room temperature or overnight at 4°C. Cells were then washed 4 times with PBS and incubated with fluorochrome-bound secondary antibodies for 1 hour at room temperature and washed 4 times with PBS. Cells were stained for 2 min with 2.5 μg/mL 4’,6-diamidino-2-phenylindole (DAPI), washed twice with PBS and overlaid with 1,4-diazabicyclo[2.2.2]octane (DABCO). Primary antibodies used include goat anti-RSV (Medix Biochemica V0601), (Milipore Sigma MABE343) and rabbit anti-PABPC1 (abcam ab21060). Secondary antibodies used include donkey anti-goat AF488 (ThermoFisher A32814) and donkey anti-rabbit AF594 (ThermoFisher A21207).

### rRNA cleavage assay

Total RNA from poly(I:C) transfected WT and *RNASEL* KO A549 cell lines was extracted using TriPure Isolation Reagent (Roche 11667165001) (according to the manufacturer’s instructions; cells grown in a monolayer option A). The RNA was quantified using the Nanodrop spectrophotometer (Thermo Fisher). Equal amount of RNA was run on a TapeStation 4150 using RNA ScreenTape Analysis (Agilent 5067-5576). Data visualization was done by TapeStation Analysis Software.

Quantification of rRNA by TapeStation analysis was performed by plating 0.05 x 10^6^ WT A549 cells in 24-well plates. Separate wells were used for cell counting by rinsing the cells with PBS, adding trypsin until all cells were lifted. Trypsin was neutralized by adding media containing FBS. The cell suspension was resuspended to break up cell clumps by pipetting up and down and counted by adding trypan blue and the Countess Automated Cell Counter (Invitrogen A49865). Live cells were counted, as determined by lack of trypan blue staining. For each RNA sample, 500 μL of TriPure Isolation Reagent containing 2 μL 100 μM 60-mer RNA spike-in was used to extract RNA according to the manufacturer’s instructions (cells grown in a monolayer option A). RNA was run on a TapeStation 4150 using RNA ScreenTape Analysis (Agilent 5067-5576) and data was visualized by TapeStation Analysis Software. Spike-in RNA and rRNA bands were quantified using Fiji and plotted with GraphPad Prism.

### Statistical testing

T-tests were 2-tailed and paired, and performed in Prism.

## Acknowledgements

We thank Guydosh and Ivanovic lab members, Alan Hinnebusch and Thomas Dever for helpful discussions. We thank Ursula Buchholz for sharing protocols and reagents for RSV titrations. James Burke for providing GADD34 A549 KO cell lines and Bernie Moss for providing WT A549 cells, PKR KO and RNASEL KO cells. We thank Antonio Gentilella for providing the LARP1 KO CRISPR-Cas9 plasmid. GADD34 expression plasmids were gifts from Shirish Shenolikar and obtained from Addgene: pXJ40 GADD34 WT (1–674 aa) (#75478), pXJ40 GADD34 (1–513 aa) (#75479), and pXJ40 GADD34 ΔPEST1 (#78096). Biorender was used to create some figure panels.

## Supplementary Figure Legends

**Figure S1.**
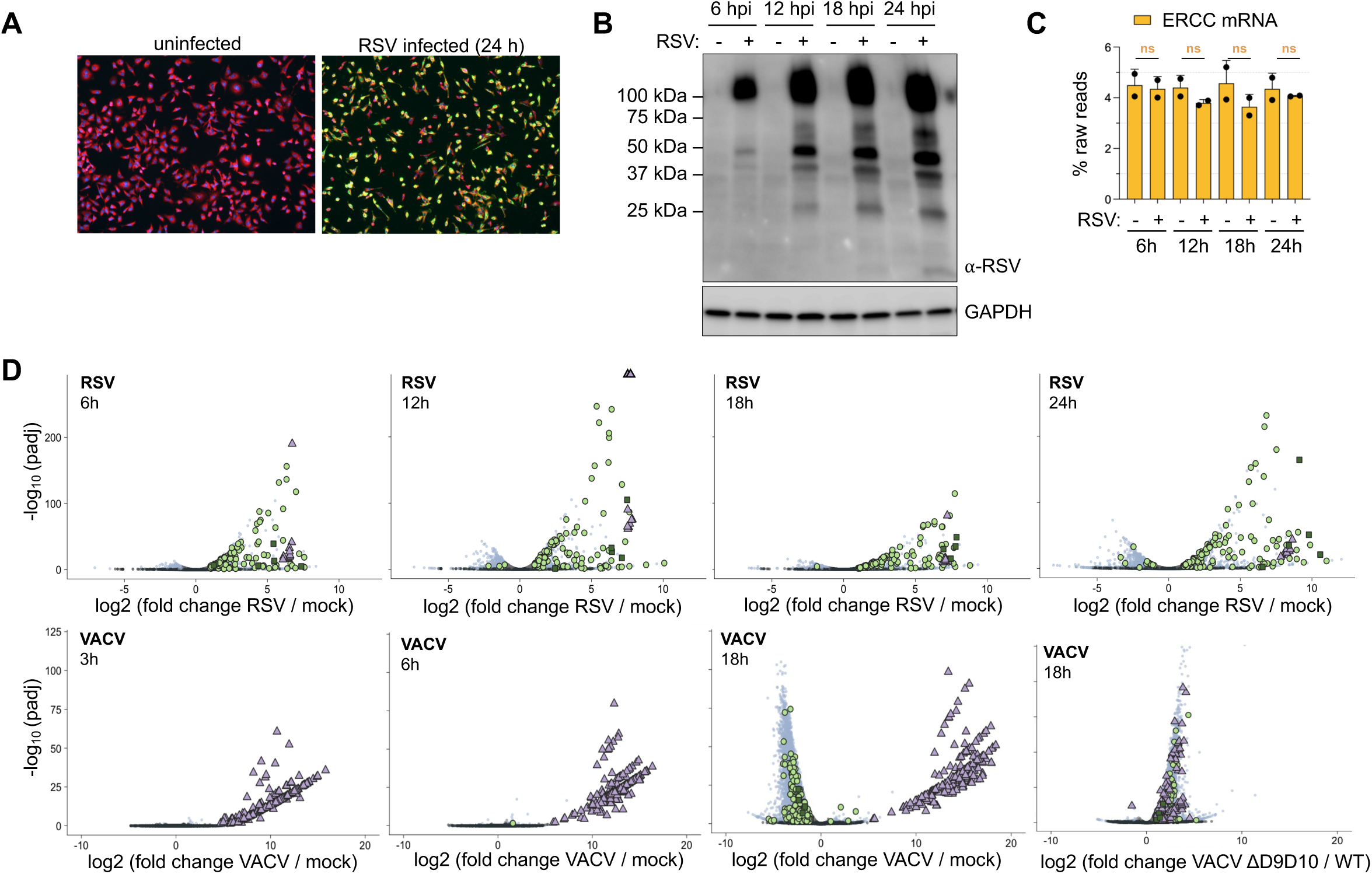
Viral, IFN and ISG transcripts are upregulated during RSV infection. (A) Immunofluorescence staining of mock- and RSV-infected cells after 24 h demonstrates a high infection rate. DAPI staining identifies nuclei (in blue), PABPC1 outlines the cytoplasm (in red) and polyclonal RSV antibody detects RSV infection (in green). (B) Western blot comparing RSV protein levels at different timepoints. RSV proteins were detected with a polyclonal RSV antibody and GAPDH serves as a loading control. (C) Quantification of the relative abundance of ERCC mRNAs, shown as a percentage of the total of raw read counts that came from host, virus and ERCC mRNAs. A paired t-test was performed to compare ERCC mRNA abundance between uninfected and infected conditions for each timepoint separately. Each datapoint indicates the value from a biological replicate with bars representing the mean and error bars indicating the standard deviation. (D) Volcano plot showing upregulation of viral mRNAs during infection. RSV infection induces a strong IFN response, while VACV infection with WT virus does not. ISG response appears to decrease due to loss of basal level of expression. Adjusted p values (padj) and log2 fold changes were calculated by DESeq2 using the Benjamini– Hochberg procedure for ERCC-normalized raw read counts. Significance cutoff was set at padj < 0.05 and log2 fold change > 1 or < -1. IFN: interferon, ISG: IFN-stimulated genes. S: significant, NC: not significant.

**Figure S2.**
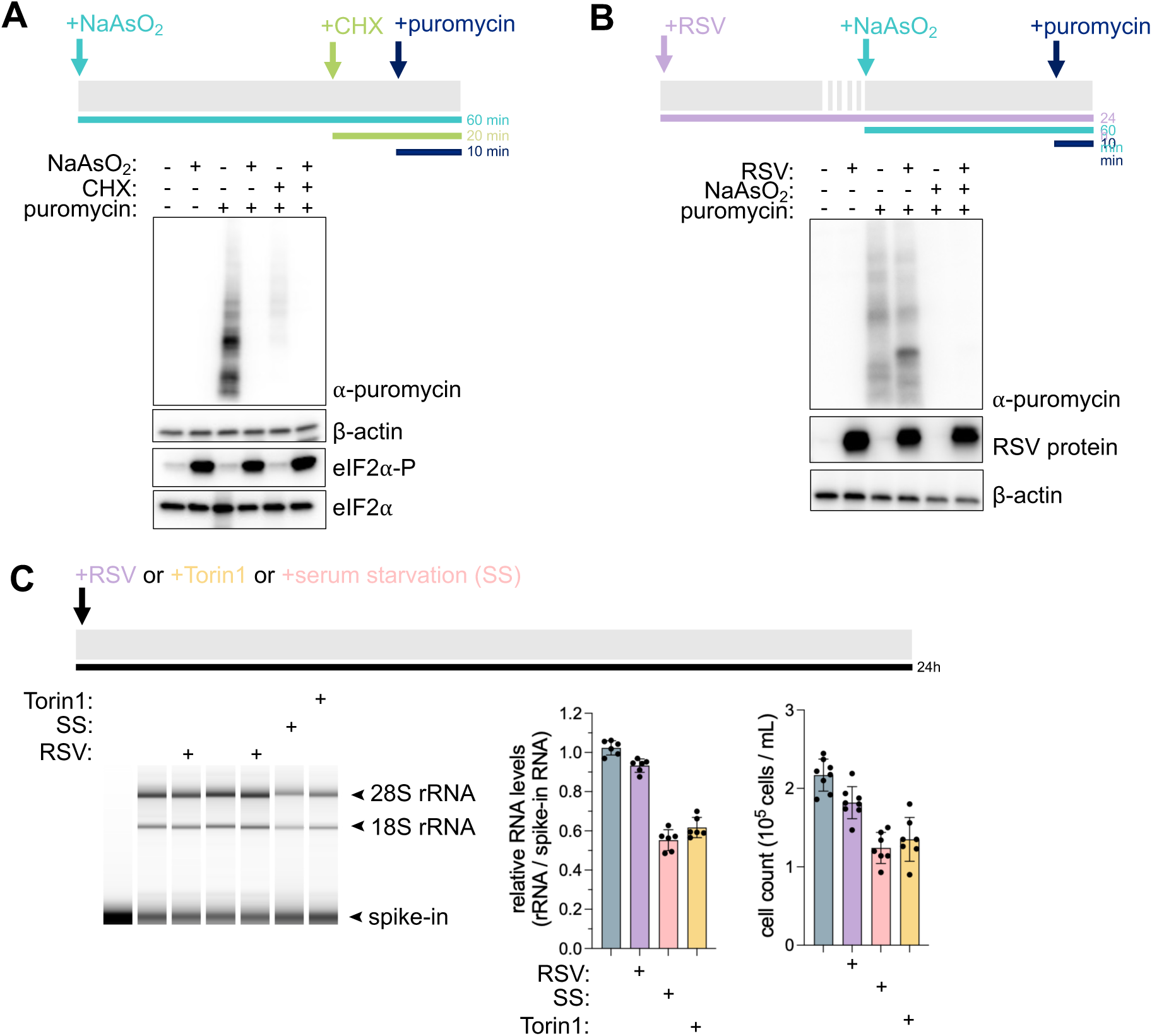
Translation and rRNA levels remain unchanged following RSV infection. (A) Western blot analysis validating the SUnSET assay by reducing translation with cycloheximide (CHX) and arsenite treatment. Cells were incubated with puromycin (10 µg/mL) for 10 minutes prior to harvest to label nascent proteins. CHX (100 µg/mL) was added 10 minutes before puromycin, as a negative control, while arsenite treatment started 50 minutes before puromycin addition. Nascent proteins were detected using an anti-puromycin antibody and β-actin served as a loading control. Detection of eIF2α and phosphorylated eIF2α-P confirmed arsenite-induced translation shutdown. Note that blot shows nascent proteins, so weights correspond to partially-completed protein. (B) SUnSET assay comparing uninfected and RSV-infected (24h, MOI 3) samples by western blot analysis. Nascent proteins were detected using an anti-puromycin antibody and RSV infection was confirmed using a polyclonal RSV antibody. Nascent protein levels are similar between uninfected and infected cells. (C) Quantitative analysis of rRNA levels by TapeStation analysis. Cells were infected with RSV (MOI 3), treated with 25 nM Torin1 or serum starved for 24 hours. Live cell counts were obtained from each well, and RNA was extracted after adding a fixed amount of 60-mer RNA spike-in per well. RNA levels for spike-in and rRNA were quantified and rRNA level per cell was computed and found to remain similar between conditions. Each datapoint indicates the value from a biological replicate with bars representing the mean and error bars indicating the standard deviation.

**Figure S3.**
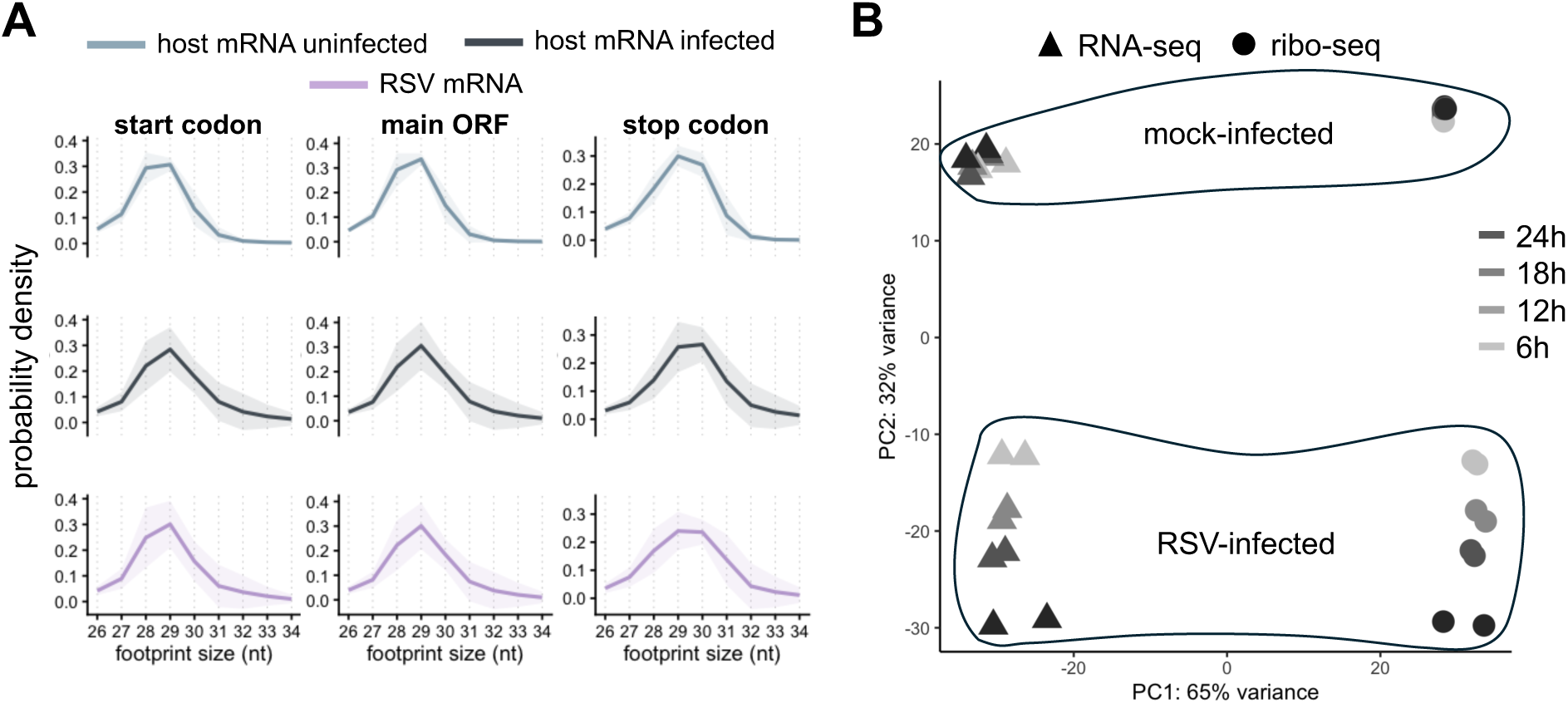
Quality control of timecourse ribo-seq and RNA-seq datasets. (A) Ribosome footprint size analysis for uninfected and infected datasets. The probability density is plotted for main ORF, start and stop codons, which serves as a quality control check for the ribosome profiling data. (B) PCA analysis for RNA-seq and ribo-seq timecourse datasets separates datasets by transcriptome and translatome (PC1) and infection duration (PC2).

**Figure S4.**
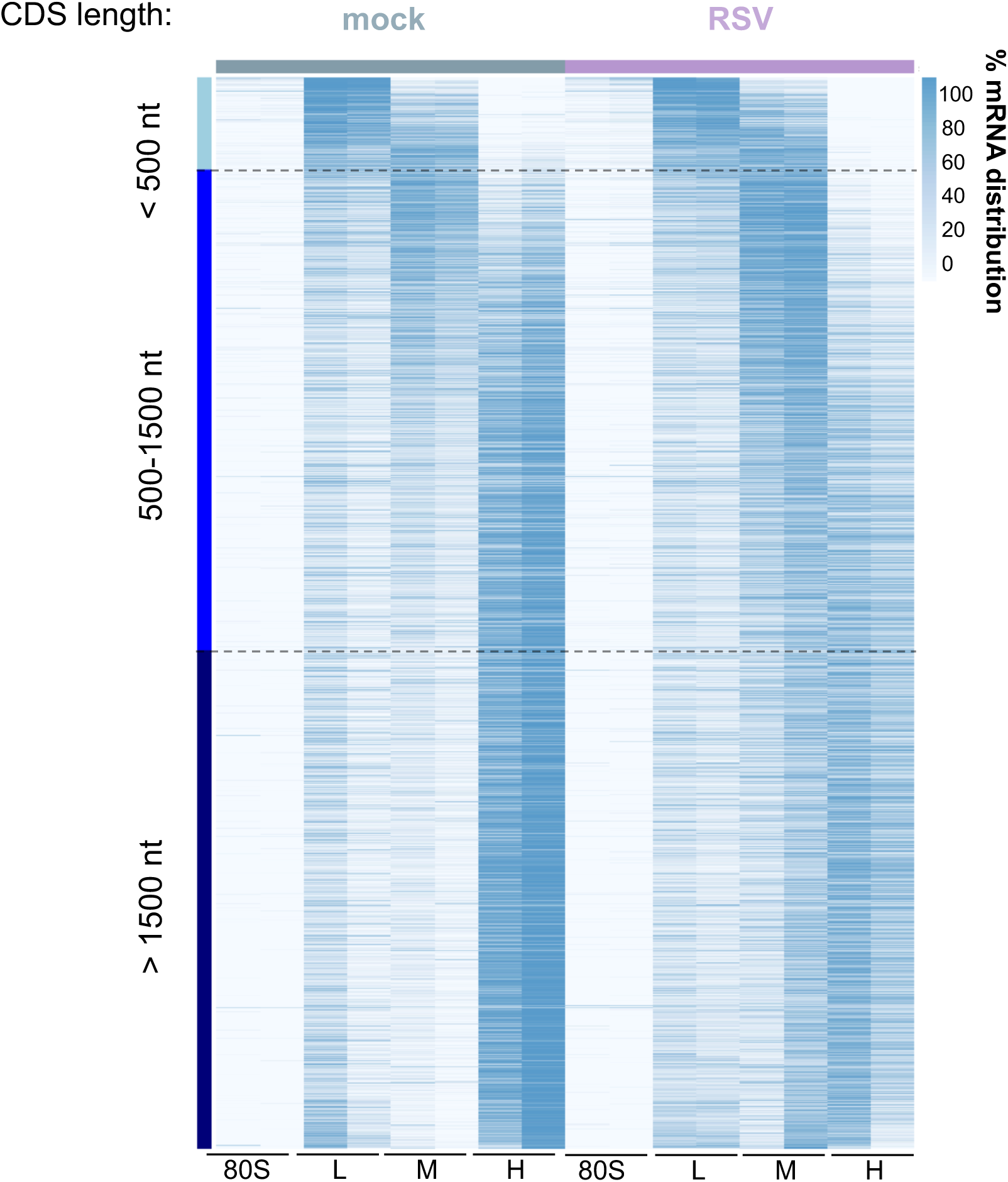
Heatmap of transcriptome-wide mRNA distribution across sucrose gradient fractions. The distribution of host transcripts along the sucrose gradient is shown as a heatmap and transcripts are sorted by main ORF length, with the shortest transcript at the top. Under uninfected conditions, transcripts with shorter ORFs are found in lighter polysome fractions while longer transcripts are in heavier polysome fractions, as expected. During infection, fewer host transcripts are found in polysomes.

**Figure S5.**
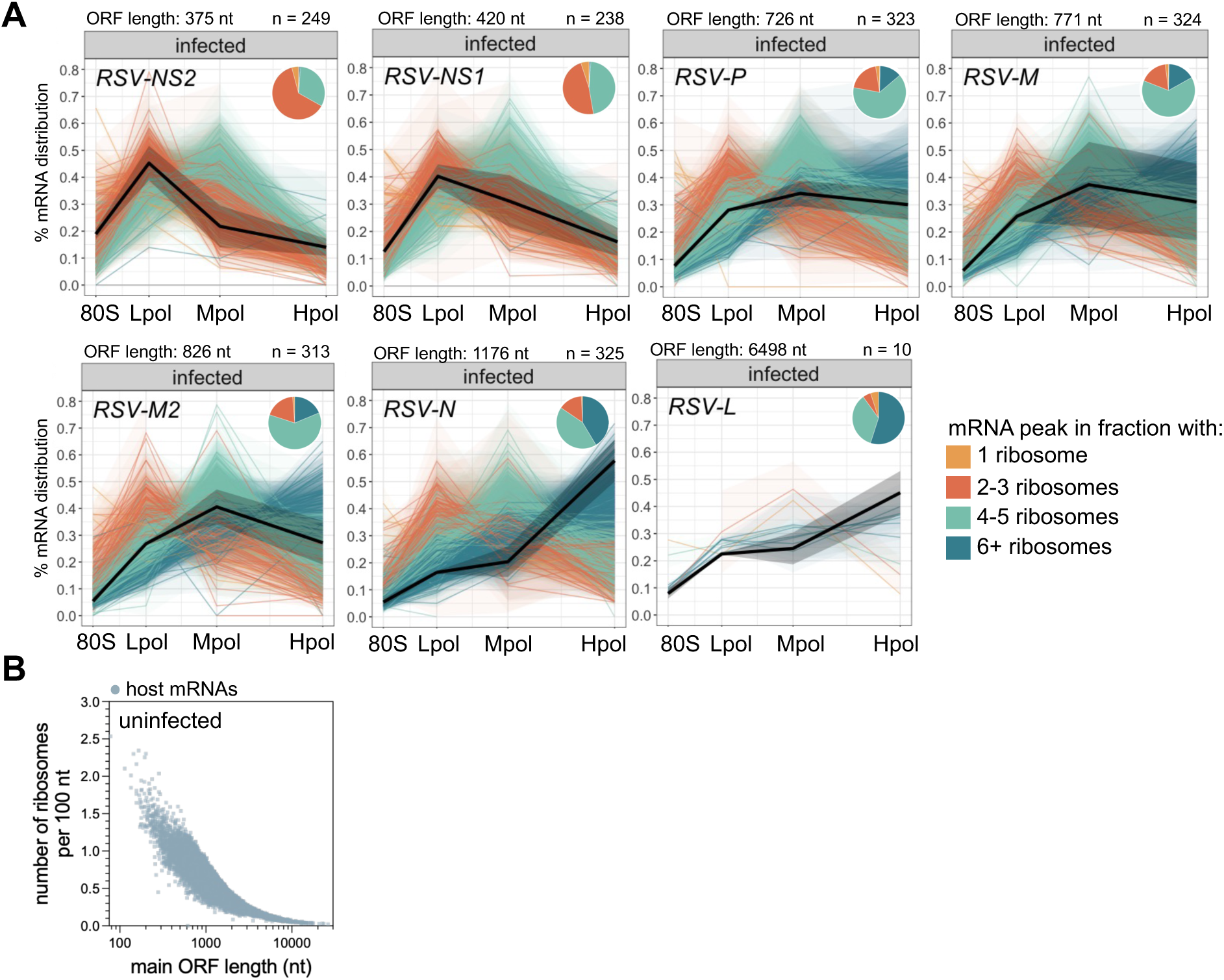
mRNA distribution of viral mRNAs and ribosome density of uninfected cells. (A) Line graphs showing the distribution of individual viral mRNAs across the sucrose gradient. The black line represents the average, and standard deviation is shown as shaded areas. Host mRNAs with similar ORF lengths (±30 nt) are colored according to their most abundant location within the gradient (80S: light orange, Lpol: dark orange, Mpol: green, Hpol: blue). Pie charts show the proportion of dominant fraction of each transcript across the gradient. The ORF length of the viral transcripts is indicated above as well as the number (n) of host transcripts included in the analysis. 80S: monosome fraction, Lpol: light polysomes, Mpol: middle polysomes and Hpol: heavy polysomes. (B) Ribosome density of individual host mRNAs in uninfected cells, calculated as number of ribosomes per 100 nt ORF length. Graph demonstrates that shorter ORFs tend to have higher ribosome density, as expected.

**Figure S6.**
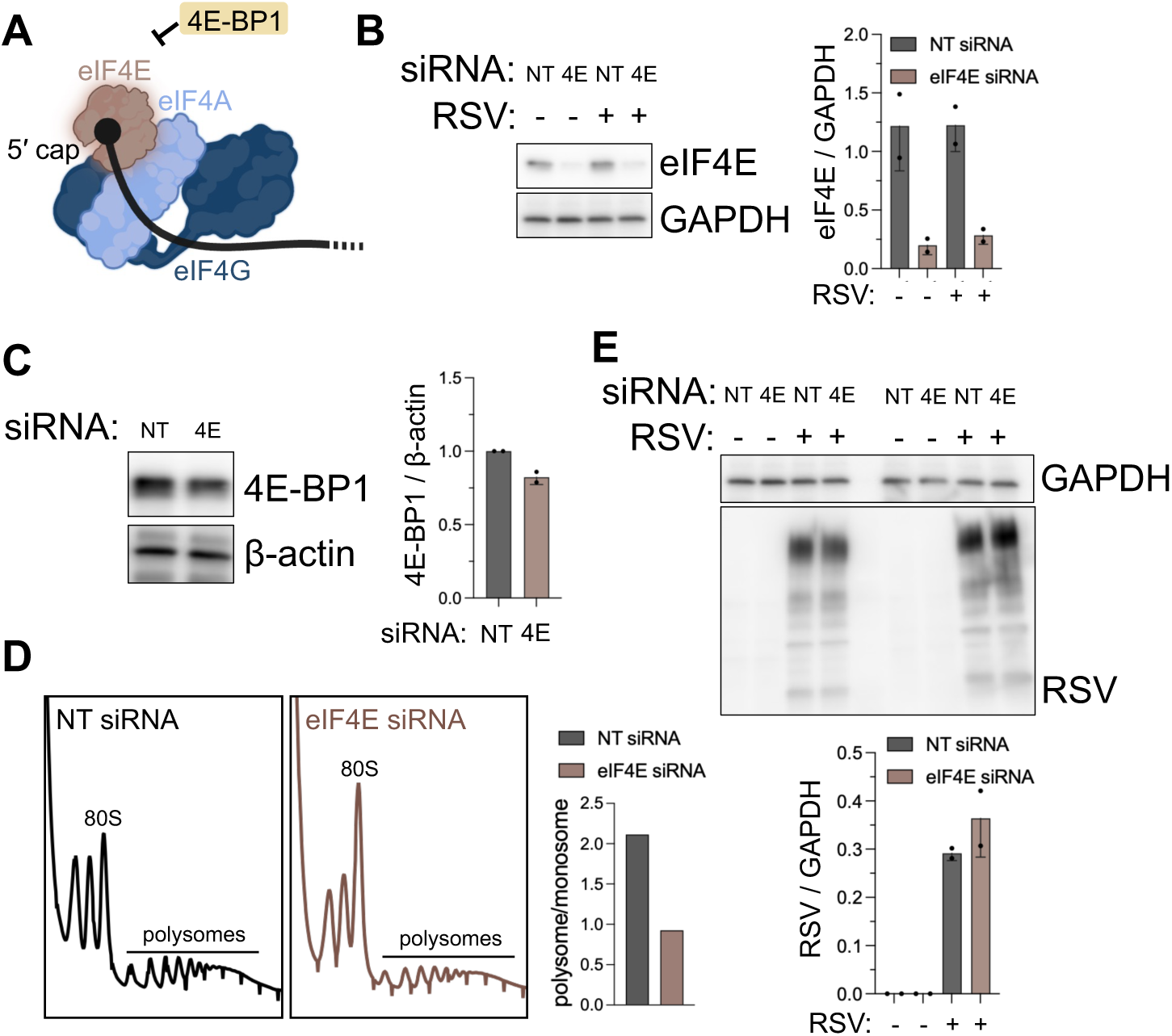
Depletion of eIF4E reduces global translation of host and viral genes. (A) Schematic overview of regulation of cap-binding protein eIF4E by 4E-BP1. (B) Western blot and quantification confirm eIF4E knockdown with GAPDH serving as a loading control. NT: non-targeting. Each datapoint indicates the value from a biological replicate with bars representing the mean and error bars indicating the standard deviation. (C) Depletion of eIF4E shows only a slight reduction in 4E-BP1 protein (shown by western blot). β-actin served as a loading control. NT: non-targeting. Each datapoint indicates the value from a biological replicate with bars representing the mean and error bars indicating the standard deviation. (D) Polysome profiling comparing non-targeting (NT) and eIF4E siRNA transfected cells shows reduction in the polysome to monosome ratio, indicating a global reduction in translation. (E) Western blot analysis shows similar levels of RSV proteins when comparing non-targeting (NT) and eIF4E siRNA transfected cells. GAPDH shown as a loading control. Each datapoint indicates the value from a biological replicate with bars representing the mean and error bars indicating the standard deviation.

**Figure S7.**
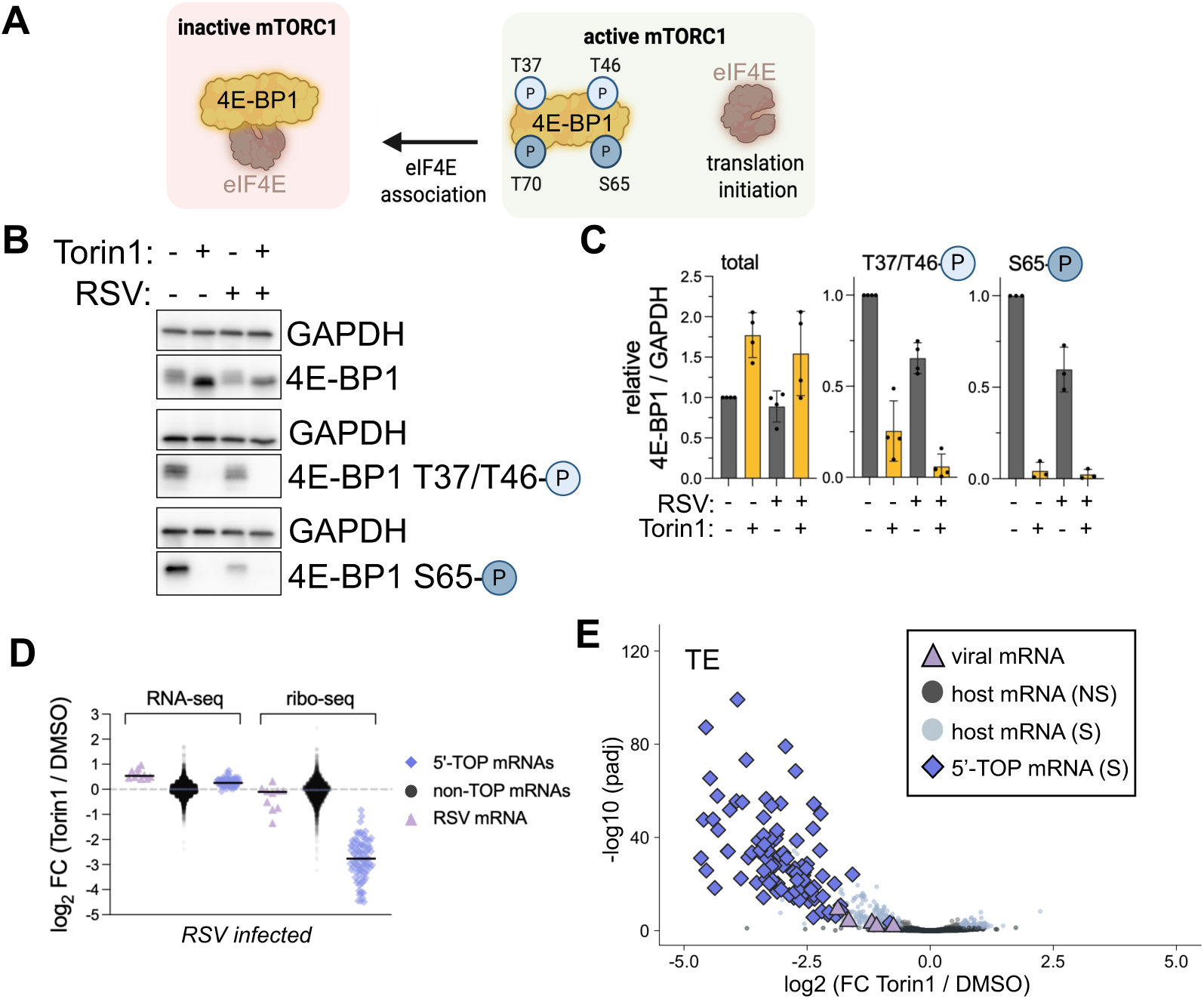
RSV translation decreases following Torin1 treatment. (A) Schematic overview of 4E-BP1 dephosphorylation which occurs following Torin1 treatment. Dephosphorylated 4E-BP1 has a high affinity for cap-binding protein eIF4E (shown on left), resulting in decreased translation initiation. (B,C) Western blot analysis of 4E-BP1 phosphorylation status following Torin1 treatment and RSV infection (B). Blot shows results obtained with three antibodies (bottom two blots are phospho-specific). Subtle shifts in band position correspond to phosphorylation status. As expected, Torin1 treatment resulted in complete dephosphorylation of 4E-BP1 (no bands in bottom two blots in lane 2 and 4). RSV infection resulted in a slight reduction in phosphorylated 4E-BP1 on both phosphorylation sites (lane 3 versus 1 of blots, see reduction in bottom two blots), as quantified in (C). Each datapoint indicates the value from a biological replicate with bars representing the mean and error bars indicating the standard deviation. (D) Differential expression analysis as computed by DESeq for all mRNAs shows the effect of Torin1 treatment in infected cells, for RNA-seq and ribo-seq datasets. The RNA abundance of 5′-TOP and RSV mRNAs was slightly increased, while the ribosome association was strongly reduced for 5′-TOP mRNAs. (E) Volcano plot showing statistical significance for changes in TE of mRNAs after Torin1 treatment of infected cells (computed by DESeq2). Data show that 5′-TOP and RSV mRNAs decrease in TE following Torin1 treatment. Adjusted p values (padj) and log2 fold changes were calculated by DESeq2 using the Benjamini–Hochberg procedure for ERCC-normalized raw read counts. Significance cutoff was set at padj < 0.05 and log2FC > 1 or < -1. S: significant, NC: not significant.

**Figure S8.**
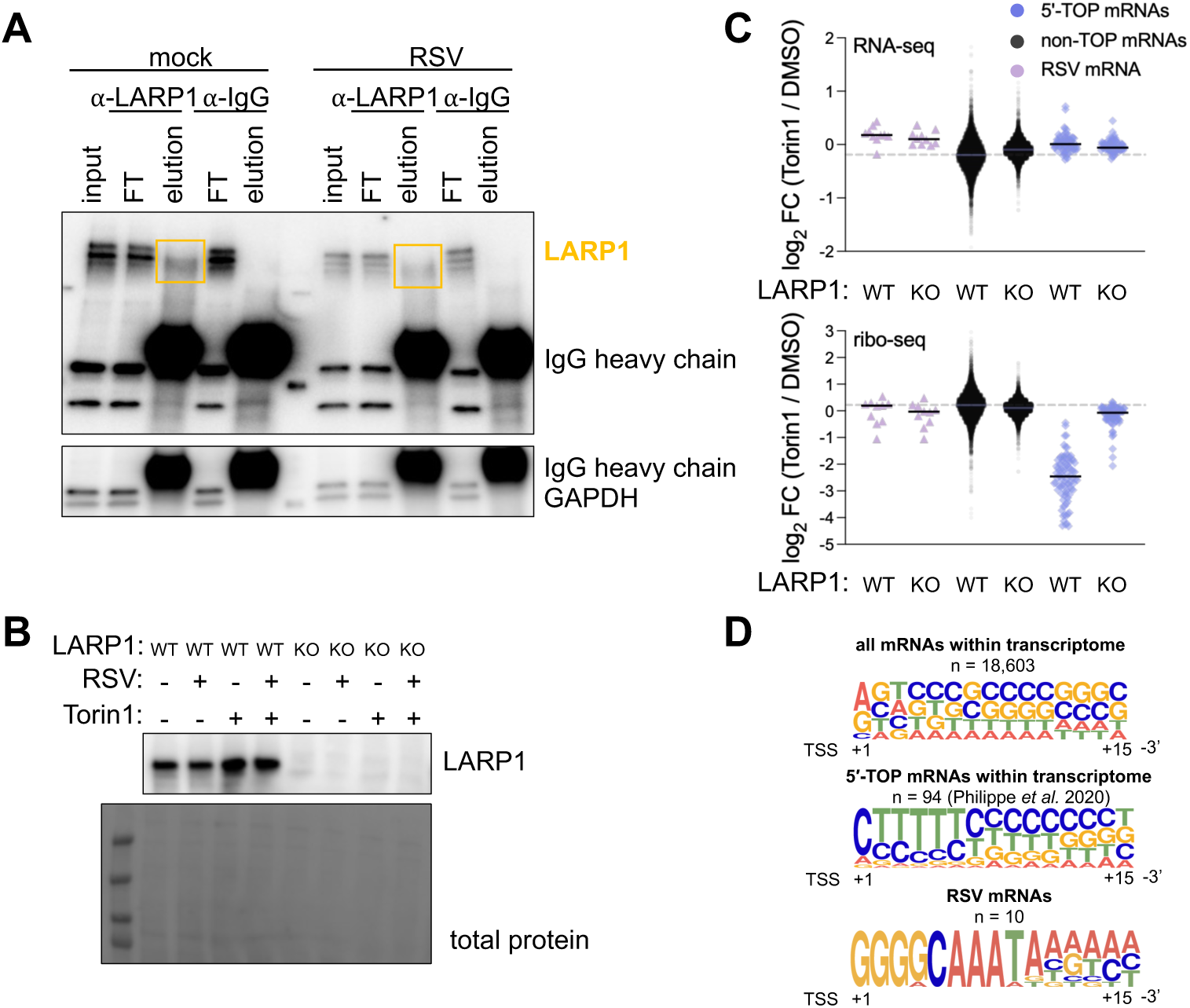
Validation of LARP1 IP and KO by comparisons between non-TOP, 5′-TOP and viral mRNAs. (A) Western blot analysis to confirm immunoprecipitation of LARP1 protein. Immunoprecipitation was performed with a LARP1 specific antibody and an IgG negative control. LARP1 was detected in input, flow-through (FT) and LARP1-IP elution samples. LARP1 protein was not detected following IgG-IP elution. GAPDH was used as a loading control and negative control (no signal in IP elution lanes). The secondary antibody used to detect primary antibodies also detected the heavy chains of the antibodies used in the IPs and are annotated on the blot (IgG heavy chain in all elution lanes). (B) Western blot confirming LARP1 KO cell lines. Total protein was detected using Ponceau staining. (C) Differential expression for all individual mRNA in WT and *LARP1* KO cells to determine the effect of Torin1 treatment in infected cells, for RNA-seq and ribo-seq datasets. The reduction in ribosome association of 5′-TOP mRNAs did not occur in absence of LARP1. (D) Motif logo of the first 15 nucleotides of an mRNA for all transcripts, 5′-TOP mRNAs and RSV mRNAs. As expected, a strong 5′ terminal oligopyrimidine tract (TOP) was observed in 5′-TOP mRNAs. This motif was absent in viral mRNAs.

**Figure S9.**
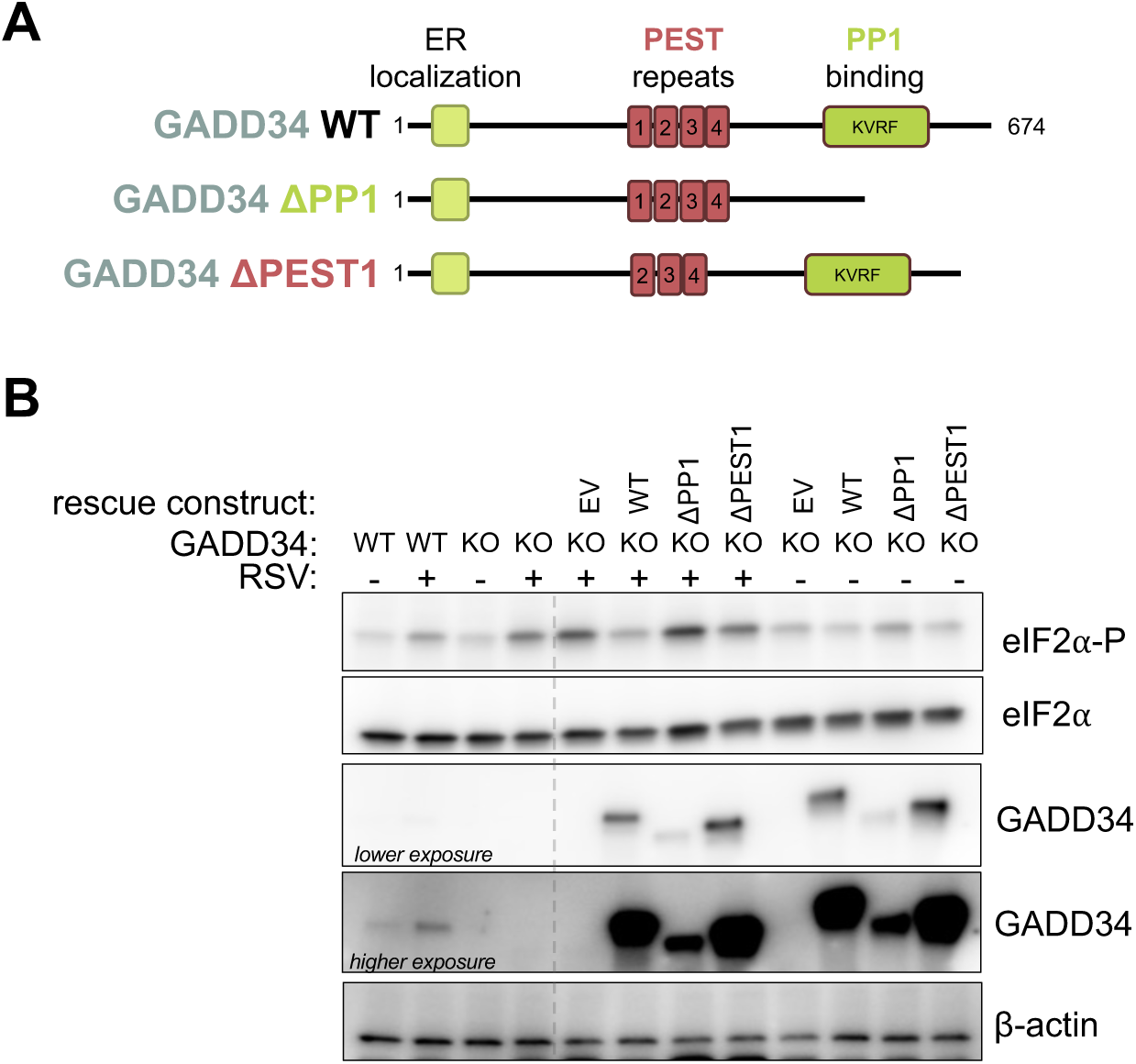
GADD34 requires phosphatase activity for eIF2α-P dephosphorylation during RSV infection. (A) Schematic representation of plasmid DNA encoding GADD34 overexpression mutants. Deletion of the PP1 domain (ΔPP1) results in deficient binding to phosphatase PP1. Deletion of the PEST1 domain (ΔPEST1) results in decreased binding to eIF2α-P. Both deletions are expected to decrease the ability of GADD34 to decrease eIF2α-P dephosphorylation. (B) Western blot analysis of rescue experiments in GADD34 KO cells. eIF2α phosphorylation levels are increased in GADD34 KO cells following RSV infection. Overexpression of GADD34 reduced eIF2α phosphorylation levels back to similar levels as WT cells. In contrast, overexpression of ΔPP1 and ΔPEST1 GADD34 mutants did not reduce eIF2α phosphorylation to background levels. β-actin and total eIF2α served as loading controls and GADD34 as KO control and plasmid overexpression control.

